# A nexus of intrinsic dynamics underlies translocase priming

**DOI:** 10.1101/2021.01.18.427065

**Authors:** Srinath Krishnamurthy, Nikolaos Eleftheriadis, Konstantina Karathanou, Jochem H. Smit, Athina G. Portaliou, Katerina E. Chatzi, Spyridoula Karamanou, Ana-Nicoleta Bondar, Giorgos Gouridis, Anastassios Economou

## Abstract

The cytoplasmic ATPase SecA and the membrane-embedded SecYEG channel assemble to form the functional Sec translocase. How this interaction primes and catalytically activates the translocase remains unclear. We now show that priming exploits a sophisticated nexus of intrinsic dynamics in SecA. Using atomistic simulations, single molecule FRET and hydrogen/deuterium exchange mass spectrometry we reveal multiple distributed dynamic islands that cross-talk with domain and quaternary motions. These dynamic elements are highly conserved and essential for function. Central to the nexus is a slender Stem through which, motions in the helicase ATPase domain of SecA biases how the preprotein binding domain rotates between open-closed clamping states. Multi-tier dynamics are enabled by an H-bonded framework covering most of the SecA structure and allowing conformational alterations with minimal energy inputs. As a result, dimerization, the channel and nucleotides select pre-existing conformations, and alter local dynamics to restrict or promote catalytic activity and clamp motions. These events prime the translocase for high affinity reception of non-folded preprotein clients. Such dynamics nexuses are likely universal and essential in multi-liganded protein machines.

## Introduction

Protein machines handle replication, transcription and unwinding of nucleic acids or folding, disaggregation, degradation and secretion of polypeptides (Avellaneda et al., 2017; Flechsig and Mikhailov, 2019; Kurakin, 2006). Such machines are commonly auto-inhibited and become activated by their partner subunits and polymeric substrates and then spend energy to remodel the latter. Their function exploits intrinsic dynamics that span multiple time regimes (Henzler-Wildman et al., 2007; Yang et al., 2014).

Unique to each protein, dynamics describe combined relative motions of protomers/subunits (hereafter ‘quaternary’), tertiary motions within a single chain (‘global’), relative ‘domain’ and ‘local’ motions (e.g. loss or displacement of secondary structure elements, hinges, loops, fluctuations of interactions between amino acids). Intrinsic dynamics usually underlie allosteric interactions however, their regulation or coupling to function remains unclear (Bhabha et al., 2015; Loutchko and Flechsig, 2020; Zhang et al., 2019).

Here we studied, the four domain DEAD box helicase member SecA that chaperones and translocates bacterial secretory polypeptides. SecA binds to the SecYEG channel in membranes to form the primed Sec translocase allosteric ensemble (Ahdash et al., 2019; Corey et al., 2019; Gouridis et al., 2013) and interacts with non-folded clients bearing signal peptides, nucleotides, lipids, chaperones and undergoes dimer to monomer transitions (De Geyter et al., 2020; Rapoport et al., 2017; Tsirigotaki et al., 2017a). Orderly, sequential ligand interactions transform the translocase from a quiescent to an active state. However, the presumably multi-tier dynamics and energetics and their link to translocation work remain elusive.

SecA, a dimeric, auto-inhibited ATPase (Fig. 1A)(Sianidis et al., 2001; Wowor et al., 2011), retains a tight ADP-stabilized state until it peripherally associates to the channel with one protomer, to form the primed translocase. This converts SecA to a 10-fold tighter preprotein binder (Fig. 1A)(Gouridis et al., 2013) and somehow prepares it for ATP hydrolysis turnovers once signal peptides and mature domain bind (Fak et al., 2004; Gouridis et al., 2009; Sianidis et al., 2001). SecA then converts to a monomeric processive motor working through mechanical strokes, Brownian ratcheting and/or by alternating channel conformations (Allen et al., 2016; Catipovic et al., 2019; Economou and Wickner, 1994; Vandenberk et al., 2019).

**Fig. 1.**
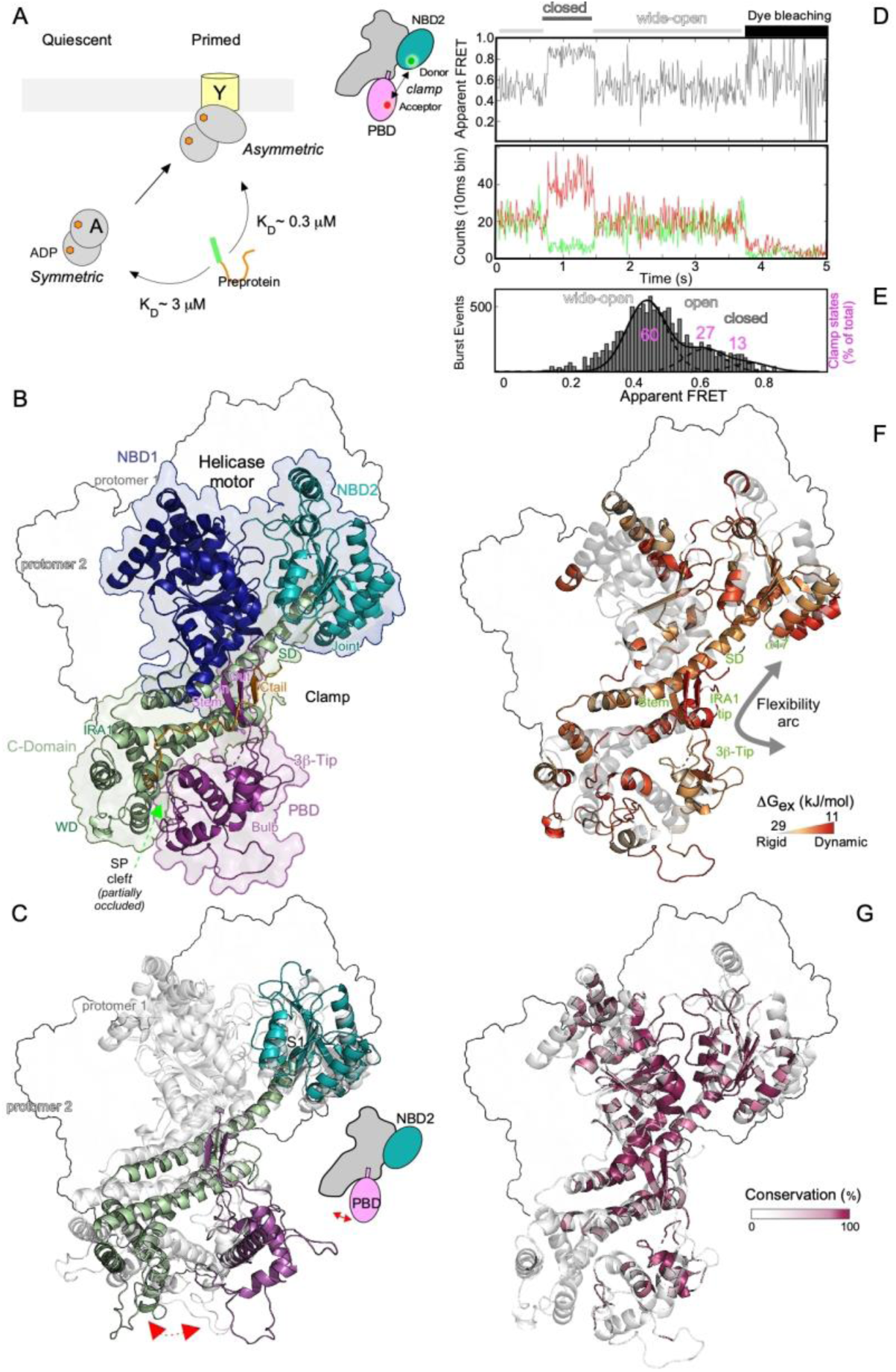
Local dynamics islands in SecA regulate clamp domain dynamics. **A.** Cytoplasmic SecA is a catalytically quiescent symmetric dimer that binds the SecYEG channel asymmetrically and forms the primed translocase holo-enzyme. Primed SecA_2_ has 10-fold higher preprotein affinity compared to quiescent SecA_2_. **B.** Domain organization of *ec*SecA_2_ (modelled after the *B.subtilis* 1M6Nin ribbon (shaded surface). ATPase motor (NBD 1 and 2: <u>N</u>ucleotide <u>b</u>inding <u>d</u>omains); PBD (<u>P</u>reprotein <u>b</u>inding <u>d</u>omain); C-domain [comprising: <u>S</u>caffold (SD), <u>W</u>ing (WD), IRA1 (Intra-molecular regulator of <u>A</u>TPase 1) and C-tail]. See also Fig. S1**. C.** MD simulations of *ec*SecA_2_,_1M6N_. Two coordinate snapshots shown from start (0 ns; grey) and end (262 ns) aligned on their NBD1 (protomer 1, coloured as in Fig. 1B and 2: contoured). **D.-E.** smFRET analysis of PBD motions using His-SecAD2, stochastically labelled with Alexa555 and Alexa647, that was either immobilized on a PEG-biotinylated-α-His antibody surface (D) or freely diffusing (50-100pM) (E). Cold His-SecAD2 (1 µM) promoted dimers (see Fig. S3A). **D.** *Top:* Representative FRET trace (out of 162) showing rare transitions between the indicated states. *Bottom:* Photon counts collected during FRET trace recording. **E.** The FRET value of every labelled SecA molecule randomly diffusing through the confocal volume (i.e. burst event, y-axis) was calculated (Apparent FRET, E*, x-axis) and plotted (left). Derived histograms (>10,000 total burst events binned in E* tranches; y axis) were fitted to a minimum of three Gaussians (Fig. S3B) representing distinct quantified clamp states (as indicated). Sum of 3 integrals= 100%; each state is a % of the total. *n=6*. **F.** ΔG_ex_ values (in kJ/mol) were calculated by PyHDX (Smit et al., 2020) from HDX-MS experiments and visualized on the dimeric *ec*SecA_2_,_1M6N_ apoprotein structure. Residues are colored on a linear scale from grey (28 kJ/mol, rigid) to red (11 kJ/mol, dynamic). Highly rigid residues (ΔG_ex_ >28 kJ/mol, transparent grey) (see also Fig. S4; Table S3). **G.** Highly conserved residues in 200 SecAs, derived from Consurf (Ben Chorin et al., 2020), coloured as indicated onto *ec*SecA_2_,_1M6N_ (represented as in C).

SecA’s helicase motor (<u>N</u>ucleotide <u>B</u>inding <u>D</u>omains 1 and 2) is fused to an ATPase suppressing C-domain and a <u>P</u>reprotein <u>B</u>inding <u>D</u>omain (PBD) (Fig. 1B; S1A-B; S1C.I). PBD, rooted via a Stem in NBD1, intrinsically rotates towards NBD2 to clamp mature domains (Bauer and Rapoport, 2009), and occupies three distinct states: Wide-open, Open and Closed (Fig. 1B; Table S1 II,III) (Ernst et al., 2018; Sardis and Economou, 2010; Vandenberk et al., 2019). Moreover, the PBD carries both the signal peptide cleft in its “Bulb” globular domain and the mature domain binding site on the narrow Stem that connects it to NBD1 (Chatzi et al., 2017)(Fig. 1B, Fig. S1B, S1C.II). When the PBD is in the Wide-open state, the dynamic C-tail of SecA folds over the signal peptide cleft and Stem regions where it binds as a pseudo-substrate guarding access to the client binding surfaces (Chatzi et al., 2017; Gelis et al., 2007).

Structures of soluble and detergent-solubilized channel/preprotein SecA states revealed their static architectures (Ma et al., 2019; Rapoport et al., 2017; Sardis and Economou, 2010) but how the underlying dynamics prime (channel binding) and activate (channel and preprotein binding) the translocase in a physiological membrane environment remain elusive. Here, we determined the intrinsic dynamics of SecA and probed how they underlie the conversion from quiescence to priming by assembly with the channel, using an integrated approach.

Fully atomistic molecular dynamics (MD) simulations and graph analysis determined ‘global’ dynamics and H-bond networks that interconnect remote regions of SecA (Karathanou and Bondar, 2019). Single molecule Förster resonance energy transfer (smFRET) reported on ‘domain’ dynamics (Gouridis et al., 2015; Kapanidis et al., 2004; Vandenberk et al., 2019) and hydrogen deuterium exchange mass spectrometry (HDX-MS) identified ‘local’ dynamics (Tsirigotaki et al., 2017b; Vadas and Burke, 2015). In both cases translocation-permitting physiological membranes and concentrations in detergent-free conditions were used.

We reveal that SecA comprises an extensive H-bond network that yields a nexus of multi-level intertwined dynamics that combine; quaternary effects from dimerization including dimer to monomer transitions, global and domain effects in each protomer, and multiple islands of local intrinsic dynamics. Within each protomer, specific islands in the helicase motor, its associated scaffold helix and the Stem conformationally crosstalk and affect the inter-conversions of the preprotein clamp between three states. ADP that occupies pre-activated SecA, tightly restricts these local dynamics but does not affect clamp motions. Dimeric, cytoplasmically diffusing SecA maintains a predominantly Wide-open clamp with weak client access. In contrast, the channel on one hand, binds to dimeric SecA, and on the other uses its carboxyterminal region to trigger re-distribution of the SecA clamp equally between the three states in the channel-bound active protomer. This structural transformation allows increased access to preprotein clients, enhanced flexibility due to relief of ADP-driven suppression of SecA local dynamics. These events thereby prepare the translocase for pre-protein mediated ADP release and activation of the enzyme for secretion work. Our data reveal how multiple ligands of a protein machine can promote step-wise priming, activation and catalysis by exploiting sophisticated and coordinated, multi-level intrinsic dynamics. The tools used here will be widely available in other membrane-embedded systems.

## Results

### Cytoplasmic SecA_2_ has restricted domain motions and a wide-open clamp

To understand how the cytoplasmic SecA_2_ is auto-inhibited but becomes primed and activated when channel-bound, we first probed its ‘global’ dynamics using simulations in bulk water. Two different, likely physiological, dimers (Gouridis et al., 2013)(Table S2), both with their clamps in Wide-open states (Fig. 1C; S2A), were used as starting structures for two independent simulations.

All four SecA domains, particularly NBD1, are extensively H-bonded (Fig. S2B) and exhibited similar dynamics irrespective of the protomer in which they belong to. Several residues participate in multiple H-bonds and/or are hubs in multi-residue H-bond pathways (Fig. S2B-D; Table S2). During 260 ns MD simulations, either of the two SecA_2_ structures only displayed minor domain motions in each of their protomers without loss of the Wide-open state (Fig. 1C; Fig. S2E). The Wide-open state is stabilized through multiple interactions that the PBD makes with the wing domain and the C-tail. This partially restricts access to the signal peptide cleft (Fig. S1C.II-III, top; Table S2).

To monitor domain motions specifically in the clamp we used our established smFRET pipeline (Fig. 1D-E; S3A)(Vandenberk et al., 2019). His-SecAD2, a single cysteinyl pair (V280C_PBD_/L464C_NBD2_) derivative, was stochastically labelled with donor and acceptor fluorophores (Vandenberk et al., 2019). The FRET state of labelled His-SecAD2 reports directly on how proximal PBD and NBD2 are and thus, on clamp motions (Fig. 1D, cartoon). Heterodimers of His-SecAD2 were generated by mixing fluorescent kinetic monomers with excess unlabeled His-SecAD2. FRET traces from surface-immobilized heterodimers were recorded as a function of time on a confocal microscope (Gouridis et al., 2015). The analysis of 162 such trajectories demonstrated that the clamp of SecA_2_ predominantly existed in a low FRET state, consistent with the ‘Wide-open’ clamp state, rarely transiting to the Open and Closed states (∼10% of the 162 traces; Fig. 1D; S3C-D)(Sardis and Economou, 2010). These data reveal that the clamp is intrinsically dynamic with individual states stable with lifetimes in the near second time regime, indicative of large domain motions (Henzler-Wildman and Kern, 2007).

Clamp states were quantified by solution smFRET. Fluorescently labelled heterodimers were monitored as they freely diffused through the confocal volume. Following pulsed interleaved excitation (Muller et al., 2005; Vandenberk et al., 2019), >10,000 photon burst events from molecules carrying both fluorophores (hence could FRET) were analyzed per experiment (Fig. 1E). 2D plots of stoichiometry vs apparent FRET efficiency were globally fitted with a mixture model of Gaussians (Fig. S3B; S3E)(Gouridis et al., 2019). The data were best fitted with 3 distributions and quantified using the area under the curve, taking the sum of all states as 100% (Fig. 1E). The clamp of SecA predominantly sampled the Wide-open state (60% of the population) and less so the Open and Closed states (27 and 13% respectively).

Collectively, these data show that SecA_2_ displays limited intrinsic domain dynamics and has its clamp predominantly in the Wide-open state. This state is maintained by several inter protomer interactions and limits access to the signal peptide cleft (Gelis et al., 2007) (Table S1; Fig. S1.C III top).

### Conserved Islands of intrinsic dynamics in SecA_2_

To define the underlying residue dynamics that define domain motions, we determined the residue level ‘local’ dynamics of SecA_2_ using HDX-MS. This technique non-invasively monitors loss or gain of backbone H-bonds, common in secondary structure, at low micromolar concentrations, near-residue resolution and in seconds timescales (Brown and Wilson, 2017; Hu et al., 2013; Skinner et al., 2012).

SecA_2_ was diluted to ∼2 µM into D_2_O buffer for various time-points. Samples were acid-quenched, protease-digested (Wowor et al., 2014) and D-uptake was determined by mass spectrometry (Fig. S4A.I). 190 peptides with high signal/noise ratio yielded ∼95% primary sequence coverage (Table S3). The D-uptake for each peptide was expressed as a percentage of its fully deuterated control (taken as 100%). D-uptake data were then processed by our in-house software PyHDX (Smit et al., 2020) to yield Gibbs free energy of exchange (ΔG_ex_, kJ mol^-1^) values for each residue. ΔG_ex_ values quantify the degree of dynamics existing within the protein backbone to which they are inversely correlated, i.e lower and higher ΔG_ex_ values represent greater and lower backbone rigidity, respectively).

SecA_2_ has several distributed regions of flexibility that together form islands of high intrinsic dynamics (Fig 1F; orange and red; Fig S4B), some of them sharply delimited against an otherwise ‘rigid’ backdrop (grey). Dynamics islands in the nucleotide binding cleft in the ATPase motor and signal peptide clefts are linked by a chain of intrinsic dynamic residues that form a ‘flexibility arc’ that lines the inner walls of the clamp (Fig 1F; grey arrow). The arc includes the 3β-tip_PBD_, peripheral loops of the PBD core, the three stranded anti-parallel β-sheet formed by the Stem (consisting of two anti-parallel β-strands and its flexible linker) and the C-tail, the tip of IRA1, the motor-associated scaffold with its kinked middle region that ultimately connects to the joint that includes α17_NBD2_s.

The nucleotide binding cleft and flexibility arc are highly conserved while the signal peptide cleft is not (Fig 1G). The local dynamics observed are likely fundamental for SecA function, and may underpin large-scale domain motions and coupling to nucleotide cycles and client binding.

### Monomeric SecA displays enhanced domain dynamics and a distributed clamp

We next determined how dimerisation affects the inherent dynamics of the SecA protomer. MD simulations of monomeric SecA revealed that PBD and NBD2 move towards each other, meet within 150 ns and form a stably Closed clamp till the end of the simulation (325 nsec). Concomitantly, C-domain sub-structures undergo domain motions (Fig. 2A; S2E-G; Movie S1). Clamp closing also exposes the signal peptide cleft (Gelis et al., 2007)(Fig. S1C.II and III, bottom; Table S1). Some residues negotiate distances of ∼2nm (Fig. S2F). During these motions PBD loses internal H-bonds (Fig. S2B). We analysed two coordinate snapshots of the Open and Closed clamp, representative conformations based on H-bond networks and centrality measurements (Fig. S2C-D, S2H). Multiple inter-domain salt bridges, dynamic H-bond clusters (mainly between PBD, scaffold and NBD2) and an extensive local hydration network lie behind clamp motions (Fig. S2I-J; Table S2). Initially, the salt-bridge of R709_WD_ with E294_PBD_ of a signal peptide cleft loop, breaks (Fig. S2H, left). Then, PBD moves towards and binds NBD2 using two prongs to form a Closed conformation (prong1: aa250-275; prong2 or 3β-tip: aa320-347; Fig. 2A; S1C.II and III, bottom; S2H, right; S2J). In this “loose Closed” conformation identified for the first time by these MD simulations, the PBD is more peripherally associated compared to the “tightly Closed” conformation seen in SecYEG:SecA:ADP.BeF_4_ crystals (Zimmer et al., 2008).

**Fig. 2.**
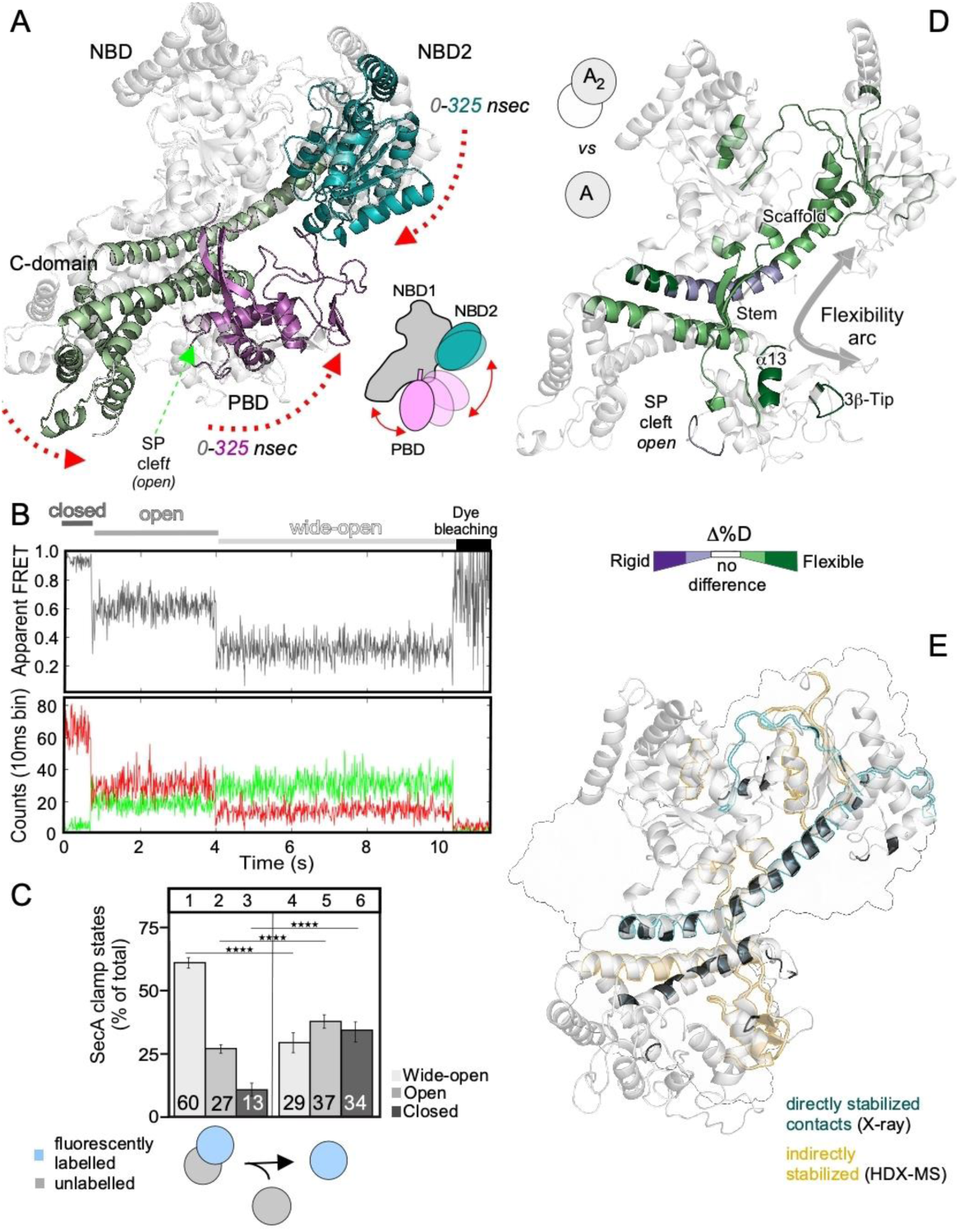
Effect of SecA. monomerization on its domain and local dynamics. **A.** Two coordinate snapshots from the start (0 ns; grey) and end (325 ns) of the MD simulations of monomeric *ec*SecA_2VDA_ aligned based on their NBD1 (grey). Domains coloured as in Fig. 1B (see also Fig. S2G-H). **B.** *Top:* Representative FRET trace (out of 84) of monomeric SecA. *Bottom:* Photon counts collected during FRET trace recording. **C.** Clamp states quantified from solution smFRET of labelled SecA_2_ with a single labelled protomer or monomers (50-100pM), as in Fig. 1E. *n≥6* biological repeats; mean ± SEM (see also Fig. S3A-B; S6A). **D.** Effect of monomerization on local dynamics of SecA. (see also Fig. S5A). D-uptake differences (light/dark hues: ΔD=10-20%/>20% respectively) between the indicated dimer/monomer end-states (control: dimer, test: monomer) are coloured on the *ec*SecA_2VDA_ structure (Top; decreased/increased dynamics: purple/green respectively; no difference: white). Flexibility arc: grey arrow. **E.** Regions of a SecA protomer stabilized in SecA_2_ either directly (involved in dimerization; *ec*SecA_1M6N_ X-ray structure; black/teal), or indirectly (from Fig. 1F; sand) as indicated.

smFRET experiments were carried out at low protein concentrations (50-100 pM) at which His-SecAD2 existed as kinetic monomers. Analysis of 84 trajectories from surface-immobilized His-SecAD2, demonstrated that the clamp of monomeric SecA freely interconverted between the Wide-open, Open, and Closed states (representative trace, Fig. 2B, S3C-D), revealed by solution smFRET to be sampled almost equally (29, 37 and 34%, respectively) (Fig. 2C, lanes 4-6; S3F; S6A I-II). Therefore, dimerization is a key extrinsic factor that suppresses the intrinsic clamp motions inherent to the monomer.

### SecA monomers display enhanced local dynamics

To monitor the local dynamics of monomeric SecA, we analysed mSecA, a fully functional derivative with reduced dimerization *K_d_* (~130 µM)(Gouridis et al., 2013), by HDX-MS. We compared mSecA to SecA_2_, focusing only on prominent differences within 5 minutes of D exchange. To ease comparison, D-uptake differences (ΔD) of a control state (Fig. 2D, upper left pictogram, top) were compared to a test one (bottom). Positive/negative values indicate regions with enhanced/suppressed dynamics (green/purple respectively), quantified as minor or major (%ΔD; 10-20% light hues; >20% dark hues).

Monomerization increased the dynamics specifically in the nucleotide binding cleft and in the flexibility arc, including most of the scaffold (Fig. 2D, green; S5A.I; Table S3). Most of the destabilized elements corresponded to regions that either directly participate in the dimer interface (Fig 2E, teal), or that are adjacent to the former, presumably reflecting allosteric effects (sand)(Fig. SIC.III top, e.g. stem, α13_PBD_ and the IRA1 second helix). The kinked middle region of the scaffold showed suppressed dynamics (purple) (Fig S4D).

### ADP rigidifies SecA_2_ and mSecA local dynamics but not clamp motion

Cytoplasmic SecA_2_ exists in a highly stable ADP state (Keramisanou et al., 2006; Sianidis et al., 2001). HDX-MS analysis demonstrated that ADP extensively stabilized multiple regions in SecA_2_ (Fig 3A, Fig S5A.II). As most of these are also stabilized in mSecA (Fig. 3B, Fig S5A.III), the ADP effects are primarily intra-protomeric.

**Fig. 3.**
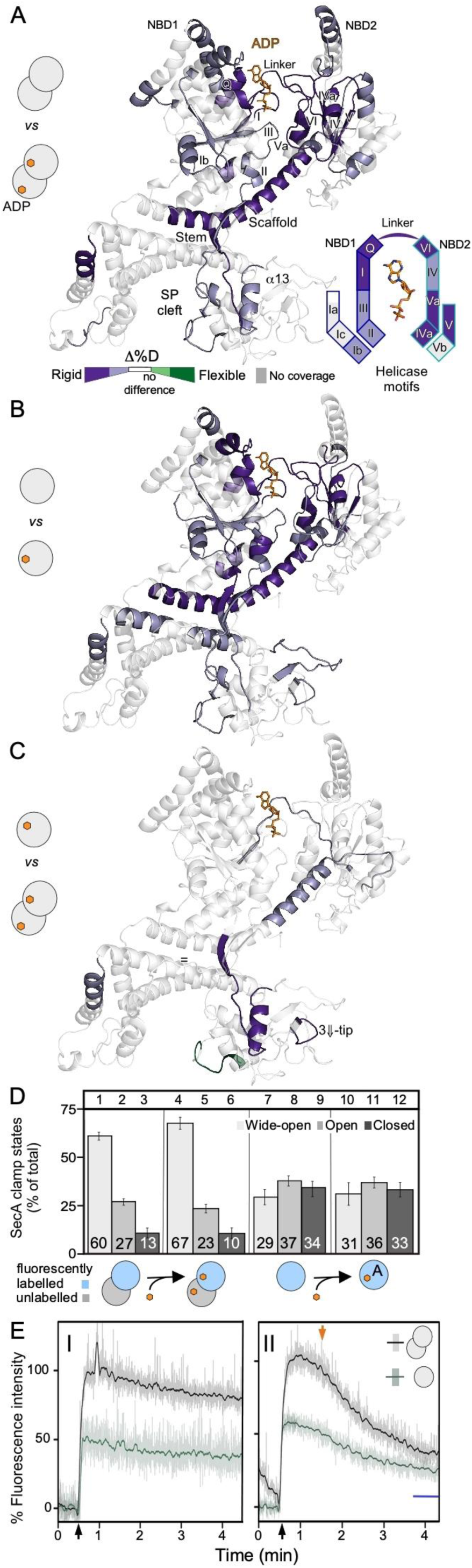
Effect of ADP on local and domain dynamics of SecA. **A.-B.** Pairwise comparisons of local dynamics determined by HDX-MS (as in Fig. 2D) upon ADP binding on SecA_2_ or mSecA. To simplify comparison, data from both proteins have been mapped on the *ec*SecA_2VDA_ “open” state (ribbon, left) and a cartoon of the helicase motifs (right). **C.** Pairwise comparison of local dynamics of ADP-bound mSecA (control) and SecA_2_ (test) to reveal the quantitative changes in dynamics that dimerization brings on top to those of ADP alone (see Fig. S5A.IV). **D.** Quantification of clamp states from solution smFRET measurements comparing apo and ADP-bound SecA dimers (as in Fig. 1E) or monomers (50-100pM; as in Fig. 1E). *n≥6* biological repeats; mean ± SEM. See also Fig. S6A. **E.** Fluorescence intensity of SecA_2_ (0.5 µM) or mSecA (1 µM), added at 30 s (black arrow) binding to MANT-ADP (1 µM) for 4.5 minutes. Raw fluorescence data (transparent lines) are superimposed on smoothened data (solid lines). The data was recorded for 4.5min normalized taking the fluorescence signal of free MANT-ADP as 0% (I) and that of SecA_2_-bound MANT-ADP as 100% (II; maximum fluorescence intensity). In II, MANT-ADP was chased with cold ADP (2 mM; added at 90 s; orange arrow). Blue line: % fluorescence intensity level of elevated ATPase mutants after ADP chase (Fig. S7A).

The nucleotide binding cleft was stabilized by multiple direct contacts that ADP makes with the helicase motifs inside the cleft and the allosteric stabilization of more peripheral ones (Fig. 3A, right panel; Fig. S1C.I), driving disorder to order transitions (Keramisanou et al., 2006). ADP binding also stabilized allosterically regions of the flexibility arc that lie several nanometers away from the nucleotide: the Scaffold (attached to the ATPase motor *in trans)* and the Stem (rooted in NBD1), that associate with each other, and the PBD that is an extension of the Stem (Fig. S1C.II). These regions provide a physical pathway for the motor to communicate with the clamp (Bondar et al., 2020). Moreover, the Stem, α13 and the 3β-tip were over-stabilized only in SecA_2_ (Fig. 3C, Fig. S5A.IV) due to pre-existing, inter-protomeric contacts in the Wide-open state that explain its stability (Fig. S1C.II-III). ADP binding stabilizes regions that were destabilized by monomerization, implying antagonism between these two processes. ADP binding drives the conformational equilibrium towards stabilization and catalytic auto-inhibition, while monomerization towards a conformationally more dynamic and, presumably, functionally activated state.

Stem dynamics are expected to regulate PBD motions and thus clamp dynamics. Remarkably, while ADP binding marginally strengthened the pre-existing contacts of the already predominant Wide-open state of SecA_2_ (Fig. 3D, compare lanes 4-6 to 1-3; Fig. S3C-D, Fig. S6B), it had no effect in monomeric SecA (compare lanes 10-12 to 7-9). In other words, the intrinsic PBD rotation occurs irrespectively of the nucleotide state of the motor. This suggested that in monomeric SecA additional factors are required to couple the local dynamics of the nucleotide binding cleft to clamp motions.

### ADP binds asymmetrically to the protomers of cytoplasmic SecA_2_

In the two SecA_2_ derivatives analyzed by MD simulations, the intra-domain H-bonding networks within each of the four domains in each protomer are similar (Fig. S2B-D). Nevertheless, they also revealed some detectable differences suggesting that despite their similarities the two protomers of the dimer retain some degree of structural asymmetry. To test if this reflects on the interaction of ADP with each protomer we monitored the binding of ADP with the environmentally sensitive fluorescent probe MANT ((2’-(or-3’)-*O*-(*N*-Methylanthraniloyl) Adenosine 5’; (Galletto et al., 2000; Karamanou et al., 2005).

MANT-ADP rapidly binds to both SecA derivatives reaching maximal intensities within 1min and remaining very stably bound for several minutes. Given its provision of two biding sites, SecA_2_ yields a ∼2-fold higher intensity than does monomeric mSecA (Fig. 3E.I). When chased with unlabeled ADP, the MANT-ADP bound to SecA_2_ is approximately halved while that of mSecA is significantly more stable and only partially exhanges (Fig. 3E.II). These data suggested that after ADP chase, SecA_2_, like mSecA, retain a single MANT-ADP tightly bound. In contrast, SecA_2_ using mutants with elevated ATPase activity, thus reduced ADP affinity, complete released the bound MANT-ADP (Fig. 3E.II, blue line; Fig. S7A).

We concluded that ADP binds to SecA_2_ asymmetrically.

### The Stem is a central checkpoint of allosteric networks

The widespread responses to monomerization or ADP binding implied the existence of extensive allosteric networks across the protein. We probed them, by graph analysis of the H-bonding networks derived from MD simulations. A large extended surface (Fig. 4A) involving many SecA residues (∼67%), are inter-connected via a dynamic water-mediated H-bond network. The Stem (Fig. 4A, dashed lines) appears to be a critical linchpin, providing a narrow passage through which the ATPase motor communicates with the PBD. To better understand its role, we probed its structural contributions further.

**Fig. 4.**
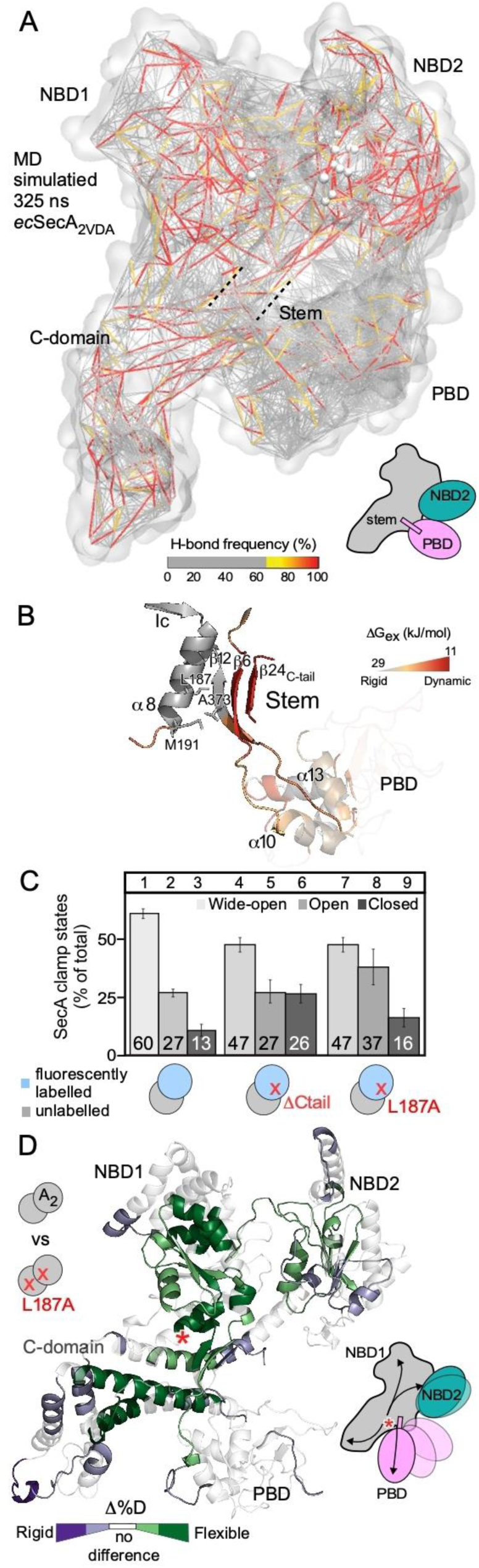
The Stem regulates the intrinsic dynamics in SecA. **A.** An H-bond protein-water network is shown for the “loose Closed” (325ns) MD simulation of *ec*SecA_2VDA_. Each line of the network represents one-water-mediated H-bond bridge in the protein-water network and is colour-coded based on frequency of appearance in the simulation. **B.** The Stem-α8 interface structure and dynamics. The Stem (β12, β6 and β24) links to α10 and α13 of the PBD. α8 is an extension of helicase motif Ic. ΔG_ex_ values of the Stem region (from Fig 1F) coloured as indicated. **C.** Quantification of clamp states from solution smFRET comparing apoSecA_2_ with SecA(ΔCtail)_2,_ and SecA(L187A)_2_ (as in Fig. 2C). *n≥6* biological repeats; mean ± SEM. **D.** Left: Pairwise comparison of local dynamics of SecA_2_ (control) against SecA(L187A)_2_ (test) (as in Fig. 2D; single protomer shown for simplicity). Right: Cartoon of SecA(L187A)_2_ (red asterisk) resulting in allosteric local effects that radiate to all SecA_2_ domains (arrows) and affect clamp motions.

The Stem is an anti-parallel β-sheet consisting of Stem_in_ (β12) and Stem_out_ (β6) of the PBD (Fig. 4B) to which a third β strand (β24_C-tail_) associates (Hunt et al., 2002). All three β strands display enhanced dynamics but lean against the rigid α8_NBD1_. While not essential for function or dimerization (Karamanou et al., 2005), β24 occupies the binding site of client mature domains (Chatzi et al., 2017). Deleting the C-tail of SecA_2_ destabilized the Wide-open state and led to clamp closing (Fig. 4C, lanes 4-6, Fig. S6B.III-IV), along with increased local intrinsic dynamics in the flexibility arc (Fig. S5B). Thus, β24_C-tail_ contributes to Stem stabilization and restriction of clamp motions.

The Stem and α8_NBD1_, share a conserved hydrophobic interface, formed primarily by α8_NBD1_ residues L187 and M191 and their juxtaposed A373 from the Stem_in,PBD_ (Fig. 4B). While the Stem as a whole is highly dynamic, the backbone of residues of β12 of Stem_in_, that participate in the Stem/α8 interface, including that of A373, are rigid (Fig. 4B). We hypothesized that this hydrophobic interface might be important for the Stem to regulate local and clamp dynamics of SecA. To test this, we weakened the hydrophobic and bulk contribution of L187 by mutating it to alanine, so as to externally affect the Stem without internally affecting the Stem β-strands. SecA_2_(L187A) is functionally active and binds preproteins with a similar affinity to SecA_2_ (Fig. S7B-C). SecA_2_(L187A) showed partial loss of the Wide-open state and consequently clamp closing (Fig. 3C, lanes 4-6, Fig. S6B.V-VI).

Remarkably, the minor mutation in L187A (Fig. 4D, red asterisk; Fig. S5C) resulted in widespread allosteric responses, mostly increased local dynamics, radiating to almost all regions of SecA_2_ (Fig. 4D).

We concluded that the Stem exploits the α8/C-tail hydrophobic interactions to directly regulate both far-reaching intrinsic dynamics networks across SecA and clamp motions (Fig. 4D, right).

### Channel-binding re-distributes the clamp in active protomers of SecA_2_

Having revealed the Stem-controlled nexus of intrinsic dynamics that extends through most of the backbone of SecA, we set out to probe if it is influenced by channel binding to yield a translocase primed for preprotein secretion.

The channel carried in inverted inner membrane vesicles (IMVs), was added at stoichiometric excess over SecA and its effect on clamp dynamics was probed. Channel binding shifts clamp equilibria from the Wide-open to Open/Closed states compared to freely diffusing SecA_2_ (compare Fig. 5A lanes 4-6 to 1-3; S6C.I-II) but not when it is mutated for interaction with SecA_2_ (SecY_M15_; Fig. 5A, lanes 7-9, Fig. S6C.III)(Karamanou et al., 2008; Matsumoto et al., 2000). Therefore, the observed alteration of clamp equilibria caused by IMVs is specific to the channel. SecY_M15_ is a conditional mutant that carries a substitution in the carboxy-terminal C-tail of SecY (Karamanou et al., 2008; Matsumoto et al., 2000), The SecY C-tail has not been crystallographically resolved but is apposed proximally to SecA, in the SecYEG:SecA complex (Fig. S7D-E) (Zimmer et al., 2008) and binds SecA directly in a peptide array (Karamanou et al., 2008). These data suggested that a functional C-tail is required for the channel to trigger clamp closing in SecA. It is important to note that this apparently occurs at a late stage of the SecA_2_-channel interaction after SecA_2_ has docked since, the SecY_M15_ channel interacts with and renders SecA a high affinity preprotein receptor (Fig. S7C) and affects it conformationally, similarly to the wild-type channel (Fig. S5E, see below).

**Fig. 5.**
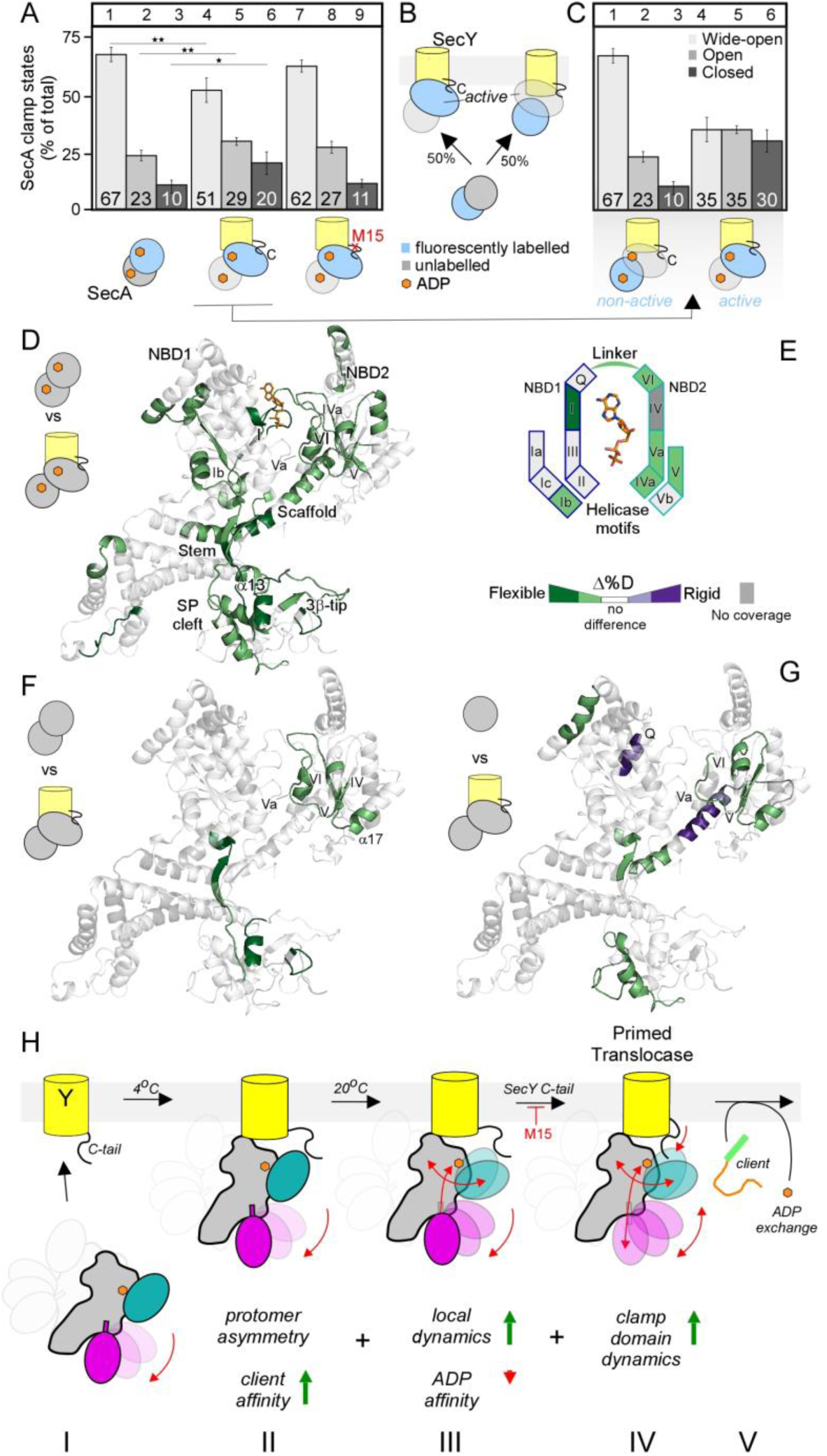
Effect of channel binding on local and domain dynamics of SecA. **A.** Clamp states of SecYEG-IMVs:SecA_2_ determined by solution smFRET (as in Fig. 2C). Fluorescence values in lanes 10-12 and 13-15 (grey gradient highlight) add up to 100% of those of the two protomers (see text for details). M15: cold-sensitive SecY derivative (Karamanou et al., 2008). *n≥3* biological repeats; mean ± SEM. **B.** Stochastic binding of SecA_2_ (with a single labelled protomer; as indicated) onto SecYEG leads to two, equally distributed, labelled or unlabelled, active SecA protomer (oval) populations. **C.** Clamp states calculated for the non-active and active protomer in channel-bound SecA_2_ (related to Fig 5A lane 4-6) **D-E.** Effect of channel binding on the dynamics of SecA_2_:ADP. D-uptake differences of channel:SecA_2_:ADP compared to SecA_2_:ADP are coloured onto the structure (D) and onto a cartoon map of the helicase motifs (E) (as in Fig 2D). **F-G.** The dynamics of channel:SecA_2_ are compared to either those of SecA_2_ (F) or of mSecA (G) apoproteins. See also Fig. S5. **H.** Model of translocase priming upon channel binding to SecA (see text for details). Asymmetric binding of soluble SecA_2_:ADP with a predominantly Wide-open clamp to SecYEG is temperature-independent and enhances local dynamics in SecA and loosening of the ADP cleft without ADP loss. >20°C, via its C-tail SecY increases clamp dynamics to the bound SecA.

This channel-induced loss of the Wide-open clamp state must be essential for translocase function. At non-permissive conditions, cells carrying SecY_M15_EG are non-viable, as the catalytic activity of the translocase was severely compromised (Fig. S7F-H)(Karamanou et al., 2008).

To properly quantify the extent of the channel-driven clamp redistribution and assign this effect to a specific SecA protomer, the asymmetric nature of SecA_2_ was considered (Fig. 3E; (Gouridis et al., 2013). SecA_2_ in solution has two functionally and structurally comparable but asymmetric protomers (Fig. 3E; Fig. 5B, bottom double circles) (Gouridis et al., 2009) and binds stochastically to the channel with only one of them that becomes the only nanomolar affinity preprotein receptor (Fig. 5B, top, ovals). The “active” protomer will also recognize the functional short C-tail of the wild-type channel but not the non-functional one of SecY_M15._ Since only one protomer in SecA_2_ is fluorescent (cyan), only half of the total measured fluorescence reflects channel-induced loss of the Wide-open clamp state in active protomers (Fig. 5A, lanes 4-6). Clamp closing exposes the signal peptide cleft for high affinity preprotein binding. The other half of the total fluorescence comes from the inactive protomers that do not contact the channel (Fig. 5B, top, circles) (Gouridis et al., 2013). These are expected to retain the domain dynamics of soluble SecA_2_ consistent with the Wide-open clamp (Fig. 5C, lanes 1-3). Subtracting the distribution of the non-active protomers from the total, yielded the distribution of the active protomer (Fig. 5C, lanes 4-6). These results revealed that the channel-primed, active protomer of SecA_2_ has its clamp almost equally distributed in three states, reminiscent of the PBD distribution in monomeric SecA (Fig. 2C, lanes 4-6).

We concluded that channel binding shifts the clamp equilibrium away from the Wide-open state and that this conformational motion is essential for translocase function.

### Channel binding to SecA_2_:ADP allosterically affects motor and Stem dynamics

We next monitored the local dynamics of the physiological SecA_2_:ADP upon channel binding. For this we developed a methodology in near-native, detergent-free conditions using IMVs (Fig. S4A.II), ensuring all available SecA_2_ was channel-bound (channel at 1.5 molar excess over SecA; 40-fold >K_d_). Peripherally bound SecA was pepsinized, IMVs removed and peptides analyzed (81% coverage; Table S3), under identical conditions to those for soluble SecA. HDX-MS averages out D-uptake values from the two asymmetric protomers of SecYEG:SecA_2_:ADP.

Channel binding resulted in increased dynamics across multiple SecA_2_:ADP regions (Fig. 5D; S5D). The most characteristic effect was the partial reversal of the extensive ADP-driven stability that had been observed in soluble SecA_2_:ADP (Fig. 3B). This included enhanced dynamics in most helicase motifs of NBD2 and only in motifs I and Ib of NBD1 (Fig. 5D and E). Despite the increased dynamics observed in many islands, dynamics of the Q motif that anchors the adenosine ring of ADP (Fig S1C.V), were unaltered, providing direct evidence that ADP remained bound. Corroborating this observation, no significant channel-driven release of MANT-ADP was detectable (Fig. S7I-J). Increased dynamics predominantly in NBD2 and in the NBD1/NBD2 linker suggested that channel binding caused NBD2 to dissociate from NBD1 (Fig. 5E). These dynamics are consistent with a helicase motor being primed by the channel for, but not yet performing, ATP catalysis (Keramisanou et al., 2006; Sianidis et al., 2001).

Elevated dynamics in the motor were directly transferred to the Scaffold, Stem and most regions of the PBD (α13, 3β-tip and the signal peptide binding cleft). Stem and PBD elevated dynamics are coincident with the enhanced clamp mobility (Fig. 5C, lanes 4-6).

To determine whether channel binding causes direct allosteric effects additionally to relieving those of ADP, we monitored the interactions of the SecA_2_ apoprotein with the channel. Channel binding enhanced dynamics in two regions of SecA_2_ (Fig. 5F; S5D): NBD2 (motifs IV, V/Va, VI, α17 to which PBD binds in the closed state) and PBD (3β-tip, α13_PBD_, Stem_in_ and C-tail). This further corroborated previous observations that channel binding results in NBD2 dynamics and liberates the clamp from the dimerization-imposed Wide-open state (Fig. 2C; S1C.III).

While the PBD distribution in the channel-bound active SecA protomer is reminiscent of that of monomeric SecA in solution, local dynamics of channel-bound SecA are nevertheless distinct from those of either free (Fig. 5G) or channel-bound (Fig. S5F) monomeric SecA, demonstrating that SecA remains dimeric upon binding the channel.

Channel-induced dynamics in SecA, prime but do not activate the motor for subsequent nucleotide cycling, while the liberated clamp prepares the translocase for binding of preprotein clients.

## Discussion

Translocase function is driven by multi-level intra-and inter-molecular intrinsic dynamics. An excessively H-bonded framework of the SecA monomer comprising flexible, distributed islands and domain motions yields a conformational repertoire of soft motion modes (Meireles et al., 2011; Zhang et al., 2020). These elements are controlled by external modulators that select pre-existing attainable conformations and contribute their own dynamics: the second SecA protomer, nucleotides, the channel and secretory clients. Such dynamics collectively support a conceptual departure from views of major ligand-biased enzyme motions between fixed start/end states, to one of subtly-balanced co-existing equilibria of dynamically interconverting states. Such mechanisms are likely generic in multi-liganded protein machines.

We dissected this dynamic landscape in depth using a multi-pronged approach under similar conditions, on physiological membranes and conditions in the absence of detergents. We linked specific intrinsic dynamics to translocase functional outcomes, i.e. the quiescent and the primed state. This pipeline sets the foundations for future studies of the translocase and other complex motors with dynamic clients, diffusing in solution or membrane-bound (Jang et al., 2019; Ramirez-Sarmiento and Komives, 2018).

Local and domain dynamics in SecA co-exist in multi-state equilibria facilitated by extensive H-bonding (Fig. 4A; S2; Table S2). Dynamics islands are sharply delimited (e.g. a few helical turns, half a β strand) and have escaped previous structural detection. They may act as either “on-off” switches (e.g. the ADP binding Walker A/motif I) or “rheostats” emanating gradients of dynamics to adjacent regions (e.g. the Stem to the Bulb). Different ligands target dynamics islands/domains differently and shift conformational equilibria; e.g. ADP affects dynamics islands in the helicase motor and scaffold but less so clamp motions.

All of the above establish an “intrinsic dynamics nexus” with distinct features at the heart of translocase catalysis. a. As the nexus exploits the pre-existing dynamics of monomeric SecA, minor free energy changes suffice for ligands to change enzyme states. We presume that this is also why ligand effects can be so easily recapitulated by point mutations that mimic signal peptide effects (Prl; (Silhavy and Mitchell, 2019) or others that render the translocase temperature-sensitive (Fig. 5A) (Ito et al., 1983; Karamanou et al., 2008; Pogliano and Beckwith, 1993). b. The multiplicity of dynamics nodes secures that SecA responds to multiple ligands, each incrementally changing its dynamics. Presumably, this also allows in the next steps of the translocation reaction the coupling of nucleotide cycling in the motor to client cycling on-off SecA and their threading through the channel. c. Long-range effects are transmitted nanometers away from a ligand interaction site as seen prominently with ADP.

Dimerization prevents monomeric SecA from expressing its excessive dynamics prematurely. This created a stable, cytoplasmically diffusing quiescent cytoplasmic state with a mainly Wide-open clamp and reduced client affinity (Fig. 1C-D; 5H.I). The two protomers of the same dimer display dynamics differences between them (Fig. S2) and therefore protomer asymmetry may predispose them to stochastic SecY binding. Channel-binding partially relieves these suppressed dynamics in the active protomer to which it binds and weakens the protomer-protomer interface while retaining dimerization (Fig. 5H.III). This allows a free tri-state Clamp distribution in the active protomer (Fig. 5.IV) and prepares the motor for ADP release (Fig. 5A-C; Fig. 5H.V). The smFRET data revealed that the Clamp in the monomeric SecA apoprotein occupies three near-isoenergetic troughs separated by activation energy barriers (Fig. 2C). Dimerization elevates the Wide-open/Open energy barrier and traps the clamp in a stable Wide-open state. In contrast, the simulated MD environment revealed a presumably energetically favoured, Open to “loose Closed” equilibrium shift in monomeric SecA (Fig. 2A; Movie S1). Crossing this energetic barrier in smFRET experiments, presumably requires the presence of both membranes and preproteins.

ADP is a top-level extrinsic regulator. It suppresses helicase motor and scaffold dynamics and hyper-stabilizes the Wide-open clamp, securing that SecA_2_ remains quiescent in the cytoplasm (Fig. 3A) and even on the channel, prior to client arrival. The main consequence of channel-induced priming is to reverse these effects to allow the free intrinsic motion of the clamp and loosen the ADP-induced restricted dynamics. In a demonstration of remarkable fine-tuning and despite its acquisition of enhanced dynamics, the helicase motor, retains ADP bound as evidenced by critical helicase motifs remaining stabilized (Fig. 5D) and fluorescence assays (Fig. S7I-J) and retains the ADP-bound asymmetry of SecA_2_. This explains how the ATPase activity of SecA_2_ is not significantly stimulated upon channel binding (Karamanou et al., 2007). Excessive stimulation will only occur once the preprotein clients bind presumably by overcoming a significant energetic obstacle driving ADP release (Fig. 5H.V). In quiescent cytoplasmic SecA_2_:ADP, a stable Wide-open clamp with its associated C-tail might impede mature domain access to the binding site (Chatzi et al., 2017), fending off unwanted cytoplasmic binders.

Protein structures are selected because their scaffolds successfully mediate specific surface chemistries. Intrinsic dynamics networks may drive their further evolution (Tiwari and Reuter, 2018; Zhang et al., 2020). Our data raise the possibility that a protein may be primarily selected because of its intrinsic dynamics propensities and then adapted to specific chemistries. The structurally conserved DEAD-box superfamily helicases, to which SecA belongs, only share sequence conservation in the helicase motifs (Jarmoskaite and Russell, 2014; Linder and Jankowsky, 2011; Papanikou et al., 2007). These motifs, many of them in weak internal parallel β-sheets (Fairman-Williams et al., 2010; Keramisanou et al., 2006; Sianidis et al., 2001), are all intrinsically dynamic (Fig. 1F-G). Such dynamics are client chemistry agnostic. The ancestral helicase motor was presumably effective in reshaping the conformational states of dynamic clients, commonly nucleic acids, albeit promiscuously and inefficiently. SecA evolved to apply the ancestral helicase motor intrinsic dynamics to amino-acyl polymer chemistries. It did this by incorporating one specificity domain that reshapes dynamic non-folded polypeptides and binds signal peptides and another that brought this chemistry to the SecY channel by associating with it.

## Experimental Procedures

### Molecular dynamics simulations

We performed atomistic MD simulations of the *E. coli* SecA monomer (*ec*SecA_2VDA_), and two independent simulations of *ec*SecA dimers. In all simulations, we considered standard protonation for all titratable groups, i.e., Asp and Glu are negatively charged, Arg and Lys, positively charged, and His groups are singly protonated. Simulation systems of the proteins in aqueous solution were prepared using CHARMM-GUI(Jo and Kim, 2008; Jo et al., 2008); ions were added for charge neutrality.

To study the dynamics of the *ec*SecA_2VDA_ monomer we used a coordinate snapshot from the NMR ensemble of SecA structures (Gelis et al., 2007). *E.coli* SecA dimer models with a Wide-open PBD and a closed ATPase motor were generated by threading the structure of the SecA monomer separately onto dimers of *B. subtilis SecA* (PDB ID: 1M6N) and *M.tuberculosis* PDB ID: 1NL3_1, two of the dimeric conformations proposed as physiologically relevant (Gouridis et al., 2013), hereafter *ec*SecA_1M6N_ and *ec*SecA_1NL3_1_. The simulation systems for the SecA monomer and dimers contain in total 345,330 (*ec*SecA_2VDA_), 635,979 (*ec*SecA_1M6N_), and 666,629 atoms (*ec*SecA_1NL3_1_).

Interactions between atoms of the system were computed using the CHARMM 36 force field (Brooks et al., 1983; MacKerell Jr. et al., 1998; MacKerell Jr. et al., 2004) with TIP3P water (Jorgensen et al., 1983). All simulations were performed with NAMD (Kalé et al., 1999; Phillips et al., 2005) using a Langevin dynamics scheme (Feller et al., 1995; Martyna et al., 1994). Geometry optimization and an initial 25ps initial equilibration with velocity rescaling were performed with soft harmonic restraints; all harmonic restraints were switched off for the production runs. Equilibration was performed in the *NVT* ensemble (constant number of particles *N*, constant volume *V*, and constant temperature *T*), and all production runs in the *NPT* ensemble (constant pressure *P*) with isotropic pressure coupling. Equilibration and the first 500ps of production runs were performed with an integration step of 1fs and all remaining production runs with a multiple timestep integration scheme using 1fs for bonded forces, 2fs for short-range non-bonded, and 4fs for long-range electrostatics. We used smooth-particle mesh Ewald summation for Coulomb interactions and a switch function between 10 and 12Å for short-range real-space interactions.

### H-bond graphs and long-distance conformational coupling

To characterize protein conformational dynamics and identify H-bond paths for long-distance conformational couplings we used algorithms based on graph theory and centrality measures (Karathanou and Bondar, 2019).

Protein groups were considered as H-bonded when the distance between the hydrogen and the acceptor heavy atom, d_HA_, is ≤2.5 Å. We computed H bonds between protein sidechains, and between protein sidechains and backbone groups. From each simulation, we derived lists of H-bonded pairs and their interaction distances and constructed adjacency matrices. These are binary matrices representing the H-bond interactions between groups (=1 if there is an H-bond connection between each amino-acid pair in the protein, and 0 otherwise). Adjacency matrices allowed us compute H-bond graphs whose nodes (vertices) are H-bonding residues, and edges, H-bonds.

For simplicity, we visualize H-bond networks by drawing unique lines between Cα atoms of pairs of residues that H-bond. These lines are coloured according to the frequency, or occupancy, of H-bonding, defined as the percentage of the analysed trajectory segment during which the two residues are H-bonded.

Using graphs, we monitored nodes and their possible interactions in the network and created clusters, i.e. paths of connected nodes (Karathanou and Bondar, 2018, 2019). To find the shortest H-bonded pathways between SecA protein domains, we use the Dijkstra’s algorithm (Cormen et al., 2009). The algorithm starts with an initial (source) and end node and finds the shortest pathway between those nodes based on positive weights. Edge weight is 1 if there is an H-bond connection between each amino-acid pair in the protein during the simulation time used for analysis and 0 otherwise. A shortest path between two nodes has the least number of intermediate nodes. We obtain the most frequently visited H-bond paths by inverting the H-bond frequencies and setting them as positive weights in Dijkstra’s algorithm.

To identify groups important for connectivity within H-bond clusters, we computed the Betweenness Centrality (BC) (Freeman, 1977, 1979) and the Degree Centrality (DC) (Freeman, 1979) of each H-bonding amino acid residue. The BC of node *n* is given by the number of shortest-distance paths that link any other two nodes (*v1*, *v2*) and pass via node *n*, divided by the total number of shortest paths linking v1 and v2. BC of node *n* can be normalized by dividing its BC by the number of pairs of nodes in the graph not including *n*. The DC (Freeman, 1977, 1979) of node *n*, equals the number of edges connecting to *n*. DC of node *n* can be normalized by dividing its DC by the maximum possible edges to *n* (which is N-1, where N is the number of total nodes in the graph). For the H-bond clusters computed here, high BC values indicate H-bonding residues that are part of many H-bond paths, whereas high DC indicates high local H-bond connectivity.

Data analyses scripts were implemented in Tcl within VMD (Humphrey et al., 1996). Additional data processing was performed using MatLab (Version R2017b, MathWorks). Unless specified otherwise, average values were computed from the last 200 ns of the monomeric and last 100 ns of the dimeric SecA simulations.

### Materials

For buffers, strains, plasmids see Supplementary Table S1, S2 and S3 respectively. D_2_O (99.9%) was obtained from Euroisotop; Alexa555 and Alexa647-maleimide from Thermo Fisher Scientific; TCEP ([Tris(2-carboxyethyl)phosphine] from Carl Roth, formic acid-MS grade from Sigma Aldrich, LC-MS grade acetonitrile LC-MS grade from Merck. All other chemicals and buffers were ACS grade from Merck or Carl Roth. proPhoA signal peptide, obtained from GenScript as lyophilized powder, was dissolved in DMSO (Merck) to a final concentration of 45 mM. Protein concentration was determined using either Bradford assay (Biorad) or/and Nanodrop^TM^ spectrophotometry (Thermo Scientific) following manufactures instructions.

### Molecular cloning

Genes were cloned in the plasmid vectors listed in Supplementary Table S3. Mutations were introduced on genes via the QuickChange Site-Directed Mutagenesis protocol (Stratagene-Agilent) using the indicated templates and primers (see supplementary Table S3). Restriction enzymes and T4 DNA Ligase were purchased from Promega. For PCR mutagenesis PFU Ultra Polymerase (Stratagene) was used; for gene amplification either Expand High fidelity Polymerase (Roche) or PFU Ultra polymerase (Promega). DpnI was used to cleave the maternal methylated DNA

(Promega). Primers (Supplementary Table S4) were synthesized by Eurogentec (Belgium). All PCR-generated plasmids were sequenced (Macrogen Europe). Plasmids were stored in DH5α cells.

### Protein purification

SecA and derivatives were expressed and purified as described (Gouridis et al., 2013; Papanikolau et al., 2007). In brief, proteins were overexpressed in BL21 (DE3) cells and purified at 4°C, using home-made Cibacron-Blue resin (Sepharose^TM^ CL-6B; GE healthcare) followed by two consecutive gel filtration steps (HiLoad 26/600 Superdex 200 pg; GE healthcare), the first in buffer A (50 mM Tris-HCl, pH 8.0, 1M NaCl), the second in buffer B (50 mM Tris-HCl, pH 8.0, 50 mM NaCl), and stored in buffer C (50 mM Tris-HCl, pH 8.0, 50 mM NaCl, 50% v/v glycerol) at −20°C. The His-tagged derivatives of SecA-D2 and proPhoA were purified as previously described (Chatzi et al., 2017; Vandenberk et al., 2019) and stored in buffer C or D (buffer C with 6 M urea) respectively. All proteins were purified to >95% purity, as assessed by gel filtration chromatography and SDS-PAGE.

SecYEG-IMVs and derivatives were prepared as previously described (Lill et al., 1989; Lill et al., 1990) and stored at −80°C. SecY concentration in these preparations was determined as described (Gouridis et al., 2013). All Sec translocase components and preprotein preparations were tested in ATPase and *in vitro* preprotein translocation assays.

### Fluorescent labeling of SecA and sample preparation for smFRET and PIE

His-SecA-D2 (10 nmol) in buffer H (50 mM Tris-HCl pH 8.0, 50 mM NaCl, 0.1mM EDTA) was treated with 10 mM DTT (1 h; 4°C), diluted to 1 mL with buffer B and added immediately onto an anion exchange resin (Q resin; GE Healthcare; 0.2 mL; equilibrated in buffer H) and incubated (5 min). The resin was subsequently washed twice with buffer H (2 x 5 mL). Alexa555-maleimide (Thermo Fisher Scientific, 50 nmol and Alexa647-maleimide (Thermo Fisher Scientific, 50 nmol) were dissolved in 5 μL DMSO. The dissolved dyes were diluted in 1 mL buffer H and added onto the resin and incubated (under gentle agitation; 12 h; 4°C shielded from light). The resin was washed with 3 mL buffer H to remove excess of dyes and allowed to settle. Proteins were eluted with buffer I (600 μl; 50 mM Tris-HCl pH 8.0, 1 M NaCl, 0.1mM EDTA). Subsequently, analytical gel filtration (Superdex 200 Increase PC 10/300; GE Healthcare) was carried in buffer J (50 mM Tris-HCl pH 8.0, 50 mM NaCl, 0.01mM EDTA) while recording the absorbance at 280 nm (protein), 555 nm (Alexa555), and 645 nm (Alexa647). The labeling ratio was estimated (>80%) based of the protein absorbance and fluorescent intensities and their corresponding extinction coefficient (ε) (εSecA = 75750 cm^-1^M^-1^, εAlexa555 = 158000 cm^-1^M^-1^ and εAlexa647 = 265000 cm^-1^M^-1^)

To study the monomeric state of SecA and derivatives the concentration of the fluorescently labelled protein was kept at 50-100 pM; the dimeric state was generated upon addition of unlabelled SecA at 0.5-1.0 μM. Incubation with different partners (ADP, unlabelled SecA, SecYEG embedded in IMVs and signal peptide) was performed at 4°C for 30min.

### Single-molecule fluorescence microscopy and PIE

Single-molecule PIE experiments were performed at 20°C using the MicroTime 200 (Picoquant, Germany). Typical average laser powers were 70 μW at 532 nm and 30 μW at 640 nm. Fluorescence emitted by diffusing molecules in solution at the focus was collected by the same water objective (UPLSAPO 60x Ultra-Planapochromat, NA 1.2, Olympus), focused onto a 75 μm pinhole and separated onto two Single-photon avalanche diodes (SPAD) with appropriate spectral filtering (donor channel: 582/64 BrightLine HC (F37-082); acceptor channel: 690/70H Bandpass (F49-691); both AHF Analysentechnik).

### PIE data analysis

Analysis was performed as described (de Boer et al., 2019; Ploetz et al., 2016). Briefly, the stoichiometry S and apparent FRET efficiency E* were calculated for fluorescent bursts having at least 200 photons, to yield a two-dimensional histogram (Kapanidis et al., 2004). Uncorrected FRET efficiency E* monitors the proximity between the two fluorophores via normalization of sensitized acceptor emission to the total fluorescence of both fluorophores during green excitation. S is defined as the ratio between the overall green fluorescence intensity over the total green and red fluorescence intensity and describes the ratio of donor-to-acceptor fluorophores in the sample.

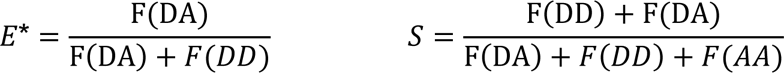

We used published procedures to identify bursts corresponding to single molecules (Eggeling et al., 2001). For this we used three parameters characterizing the burst: total of L photons with M neighbouring photons within a time interval of T microseconds. For the data presented in this study, a dual-colour burst search (Nir et al., 2006), using parameters M = 35, T = 500 μs and L = 50, was applied. Additional thresholding removed spurious changes in fluorescence intensity and selected for intense single-molecule bursts (all photons > 200 photons unless otherwise mentioned). E* and S values for each burst and thus for individual molecules were binned into a two-dimensional histogram, where we selected donor-acceptor-containing sub-populations according to their intermediate S values. The one-dimensional E* histograms were fitted with a mixture model of a variable number of Gaussian distributions (1-3). In the fitting procedure the mean and the amplitude were derived from fitting, whereas the standard deviation was fixed or allowed to vary over a small region defined from static DNA samples having attached fluorophores at specific positions (Fig S3). We used the minimum number of distributions that fitted the experimental data, in which the mean value defines the apparent FRET value (E*) and the amplitude the abundance of a conformational state.

### Confocal scanning microscopy and data analysis

To gain information on possible conformational sampling of SecA at room temperature, we used the same home-built confocal microscope as described before (Gouridis et al., 2015). Surface scanning was performed using a XYZ-piezo stage with 100×100×20 μm range (P-517-3CD with E-725.3CDA, Physik Instrumente). The detector signal was registered using a Hydra Harp 400 picosecond event timer and a module for time-correlated single photon counting (both Picoquant). The data, e.g., time traces and scanning images, were extracted using custom made software. Data were recorded with constant 532-nm excitation at an intensity of 0.5 μW. Surface immobilization was conducted using an anti-His antibody and established surface-chemistry protocols as described (Gouridis et al., 2015).

### Quantification and statistical analysis for FRET data

Statistical analysis was performed with Origin software version 2018 (OriginLab), Matlab R2014b (MathWorks). FRET histograms were fitted with a Gaussian mixture model with a restricted standard deviation (see methods section for details). The data (means and the amplitudes) correspond to mean of 3-5 repeated experiments (i.e. independent protein purification and labelled sample).

### Amide hydrogen/deuterium exchange mass spectrometry

HDX sample preparation: SecA and derivatives (see Supplementary material) were dialyzed overnight into buffer E (50 mM Tris-HCl pH 8.0, 50 mM KCl, 1 mM MgCl_2_, 1 mM DTT) and concentrated (∼100 µM) using centrifugal filters (Vivaspin 500, Sartorius). In the *apo* condition, the protein stock was diluted into aqueous buffer B at 1:5 ratio prior to dilution in D_2_O. For the ADP-bound state, SecA was incubated with 20 mM nucleotide, prior to dilution into D_2_O (2 mM final nucleotide concentration in D exchange reaction). To monitor SecA:proPhoA_1-122_ interactions, proPhoA_1-122_ (in Buffer D) was diluted in buffer E to a final concentration of 250 µM (0.2 M Urea), immediately added to SecA at 1:10 ratio (SecA: proPhoA_1-122_) and incubated for 2 minutes prior to D exchange. For the SecA:SecYEG state, IMVs were sonicated as described (Chatzi et al., 2017; Gouridis et al., 2010) and incubated with SecA at 1:1.5 (SecA:SecY) molar ratio, for 2 min on ice, prior to D exchange. To monitor how signal peptides (signal peptide) activate the translocase, the synthetic proPhoA signal peptide (Genescript; 45mM in 100 % DMSO) was diluted 30-fold into Buffer E (to obtain 1.5 mM signal peptide in 3 % DMSO), added to preincubated SecA:SecYEG (see above) at a final molar ratio of 2:3:15 (SecA:SecYEG:signal peptide) and the reaction was incubated for a further 1 minute. All mutant proteins were handled similar to the wild type ones and reactions were maintained at similar molar ratios.

#### D exchange reaction

Isotope labeling was carried out using lyophilized buffer F (50 mM Tris-HCl pH 8.0, 50 mM KCl, 1 mM MgCl_2_, 4 µM ZnSO_4_) reconstituted in 99.9% D_2_O (Euriso-top), with fresh TCEP [tris(2-carboxyethyl)phosphine] added at 2 mM. Buffer pH_read_ was adjusted to 8.0 using NaOD (Sigma). D exchange buffer was pre-incubated in a 30°C water bath, and the D exchange reaction was initiated by diluting 200 pmol of protein into D_2_O buffer F at a 1:10 ratio (final D_2_O concentration 90%). Final concentration of SecA was maintained at 4 µM in the D exchange reaction. Continuous labeling reaction was incubated for various time points (10 s, 30 s, 1 min, 2 min, 5 min, 10 min, 30 min and 48 h), primarily at 30°C. For D exchange experiments carried out at 18 °C, fewer time points were obtained (1 min, 5 min, 10 min, 30 min, 60 min).

#### Quenching

The D exchange reaction was quenched by the addition of pre-chilled quench buffer G (1.3% formic acid, 4 mM TCEP, 1 mg/mL fungal protease XII) at a 1:1 ratio (final pH of 2.5), and incubated (4°C; 2 min). In samples containing SecYEG IMVs, the reaction was centrifuged at 20000 x g for 90 s on a benchtop cooled centrifuge (Eppendorf), the supernatant containing SecA peptides was collected and immediately injected into the LC-MS system. 100 pmol of SecA was injected into a nanoACQUITY UPLC System with HDX technology (Waters, UK) coupled to a SYNAPT G2 ESI-Q-TOF mass spectrometer. For enhanced peptide coverage, SecA was digested in 2 steps, first with fungal protease XIII (Sigma) (Wowor et al., 2014) at the quench step, and subsequently online digestion on a home-packed immobilized pepsin (Sigma) cartridge (2 mm x 2 cm, Idex), at 16°C. The resulting peptides were loaded and trapped onto a VanGuard C18 Pre-column, (130 Å, 1.7 mm, 2.1 x 5 mm, Waters) at 100 mL/min for 3 min using 0.23% (v/v) formic acid. Peptides were subsequently separated on a C18 analytical column (130 A°, 1.7 mm, 1 x 100 mm, Waters) at 40 mL/min. UPLC separation (solvent A: 0.23% v/v formic acid, solvent B: 0.23% v/v formic acid in acetonitrile) was carried out using a 12 min linear gradient (5-50% solvent B). At the end, solvent B was raised to 90% for 1 min for column cleaning. Peptide trapping-desalting and separation were performed at 2°C. The MS parameters were as follows: capillary voltage 3.0 kV, sampling cone voltage 20 V, extraction cone voltage 3.6 V, source temperature 80°C, desolvation gas flow 500 L/h at 150°C. D exchange experiments were carried out in technical triplicates for most conditions (details in Table S3), and experiments were performed over multiple days to control for day to day instrument variation. Further, SecA apo data were obtained from 3 separate protein purifications (biological triplicates) and data was compared to check for any biological or technical variability. Full deuteration controls were obtained by incubating SecA in buffer F containing 6M Urea-d4 (98% D, Sigma) overnight at room temperature. D-uptake (%) was calculated using the full deuteration control D-uptake values. Deuterium/Protium back exchange values for our instrumental set up was calculated to be between 20-45% depending on peptide composition. These values are consistent with previously reported studies using similar instrumental set ups (Walters et al., 2012). The data has not been corrected for back exchange and is represented either as absolute D values or as a percent of the full deuteration control (Wales et al., 2013).

### Peptide identification and HDX data analysis

Peptide identification was carried out using 100 pmol of protein diluted in protiated buffer F. The sample was quenched as described above and analysed in the MS^E^ acquisition mode in a nanoACQUITY UPLC System with HDX technology (Waters, UK) coupled to a SYNAPT G2 ESI-Q-TOF mass spectrometer over the m/z range 100-2,000 Da. The collision energy was ramped from 15 to 35 V. Other instrument parameters were as described above. Peptide identification was performed using ProteinLynx Global Server (PLGS v3.0.1, Waters, UK) using the primary sequence of SecA or derivatives as a search template. Peptides were individually assessed for accurate identification and were only considered if they had a signal to noise ratio above 10 and a PLGS score above 7.5. Further, peptides were only considered if they appeared in 3 out of 5 replicate runs for each protein. Data analysis was carried out using DynamX 3.0 (Waters, Milford MA) software to compile and process raw mass spectral data and generate centroid values to calculate relative deuteration values. All spectra were individually inspected and manually curated to ensure accurate centroid calculations. Maximum errors between replicate runs were found to be ±0.15 Da with most errors within ±0.08 Da, thus a difference of ± 0.5 Da between peptides from different states was considered significant (Houde et al., 2011).

### HDX-MS data interpretation and visualization

Comparison between different states of SecA was carried out by considering one state as the control and the other as the test state. D-uptake values were first converted to %D values (as a percentage of 100% deuteration control). %D values of the control state were subtracted from the test state. Positive values indicated increased dynamics and negative values indicated decreased dynamics in the test state compared to the control state. Comparison between states was carried out only on the same peptide and time point obtained from different states. A significant difference in D-uptake in a peptide between two states was identified if it satisfied 2 criteria: a. > ± 0.5 Da absolute difference in deuterium exchange (Houde et al., 2011), and b. >10% difference in % D values between the 2 states. A peptide was considered different in dynamics if 1 or more time points showed significant differences in D-uptake between 2 compared states. Differences observed within 5 min of D exchange were weighted with greater importance as these time points are hypothesized to monitor the determinants of sub-second domain motions (as determined by smFRET). The peptides that showed significant differences were further classified into those with minor and major differences based on difference in % exchange between the two states; differences between 10%-20% were considered minor and differences > 20% were considered major changes.

### Binding of MANT-ADP to the Sec translocase

Binding of MANT-ADP to the ATPase motor of SecA results in a strong increase in fluorescence intensity due to the hydrophobic environment within the nucleotide binding cleft (Galletto et al., 2000). Fluorescence intensity was measured on a Cary Eclipse fluorimeter (Agilent) at fixed wavelengths λ_ex_= 356 nm and λ_em_= 450 nm. Excitation slit was at 2.5 nm and emission slit was at 5 nm. All experiments were carried out in 1 mL of buffer K (50 mM Tris-HCl pH 8, 50 mM NaCl, 1 mM MgCl_2_). MANT-ADP was maintained at 1 µM. 1 µM of SecA/SecYEG:SecA was added at t=30 sec and fluorescence intensity was monitored for 5 min. In ADP chase experiments, 2 mM of cold ADP was added at t= 90 sec and fluorescence was monitored till t= 5 min. Fluorescent measurement graphs were smoothened using cubic spline smoothening (GraphPad Prism 5).

### Miscellaneous

Structural analysis was performed and movies were generated using Pymol (https://pymol.org/) and sequence conservation was visualized onto the structure using Consurf (Ben Chorin et al., 2020). Affinity determination of SecA and/or proPhoA for the translocase, SecA ATPase activity, *in vivo* proPhoA and PhoA translocation, *in vitro* proPhoA translocation, SecA activation energy determination, *in vivo* SecA complementation were as described (Chatzi et al., 2011; Gouridis et al., 2009; Gouridis et al., 2010; Gouridis et al., 2013).

## Acknowledgements

This paper is dedicated to the memory of Yiannis Papanikolau who solved the first *E.coli* SecA structure. We are grateful to: M. Papanastasiou for preliminary HDX-MS analysis of SecA and A.Tsirigotaki, M.B.Trelle, J.T.D. Jorgensen for discussions and advice; T. Cordes for sharing software for smFRET data analysis and V. Krasnikov for advise in setting up the smFRET microscope. Research in the Economou lab was funded by grants (to AE): RiMembR (Vlaanderen Onderzoeksprojecten; #G0C6814N; FWO); MeNaGe (RUN #RUN/16/001; KU Leuven); ProFlow (FWO/F.R.S.-FNRS “Excellence of Science - EOS” programme grant #30550343); DIP-BiD (#AKUL/15/40 - G0H2116N; Hercules/FWO); PROFOUND (Protein folding/un-folding and dynamics; W002421N; WoG/FWO) and (to AE and SK): FOscil (ZKD4582 - C16/18/008; KU Leuven) and (to AE and GG): CARBS (#G0C6814N; FWO). Funds for the smFRET microscope were from ZKD4582 and Economou lab resources. Research in the lab of A-NB was supported in part by the Excellence Initiative of the German Federal and State Governments via the Freie Universität Berlin, and by allocations of computing time from the North-German Supercomputing Alliance, HLRN. GG was funded by the NWO (Veni grant 722.012.012), the Zernike Institute for Advanced Materials and the Rega Foundation postdoctoral program; SKr was a FWO [PEGASUS]² MSC fellow, JHS is a PDM/KU Leuven fellow and NE is a MSCA SoE FWO fellow. This project has received funding from the Research Foundation – Flanders (FWO) and the European Union’s Horizon 2020 research and innovation programme under the Marie Skłodowska-Curie grant agreements No 665501 and 195872.

## Author contributions

SKr purified proteins and membranes, did biochemical assays, designed and performed HDX-MS work and data analysis, performed ADP release assays and homology analysis. NE purified and labelled proteins and performed smFRET calibrations and experiments and data analysis, analysed structures and made movies. KK performed MD simulations and graph analysis of H-bond networks. JHS developed the PyHDX software and adapted FRET burst analysis for Microtime200 output data. AGP produced the HDX-MS and smFRET constructs by molecular cloning and mutagenesis. KEC performed molecular biology and *in vivo* assays. SK designed and supervised molecular biology experiments and smFRET constructs, purified proteins for smFRET, performed and supervised biochemical and biophysical assays and data analysis. ANB set up and supervised the MD simulations and performed runs and data analysis. GG adapted the smFRET pipeline to His-tagged SecA and designed the immobilized smFRET with major contributions from NE, technically supervised the smFRET work, the installation of the Microtime200 set up, performed molecular biology, biochemical and biophysical assays, analysed data. AE designed experiments, did structure and data analysis, performed biological/structural interpretation and integration of HDX-MS, smFRET and MD data. AE and SKr wrote the first draft with contributions from NE, KK, JHS, AP, ANB, KK, SK and GG. All authors reviewed and approved the final manuscript. AE and SK conceived and managed the project.

## Supplement

### Supplementary Figures

**Figure S1.**
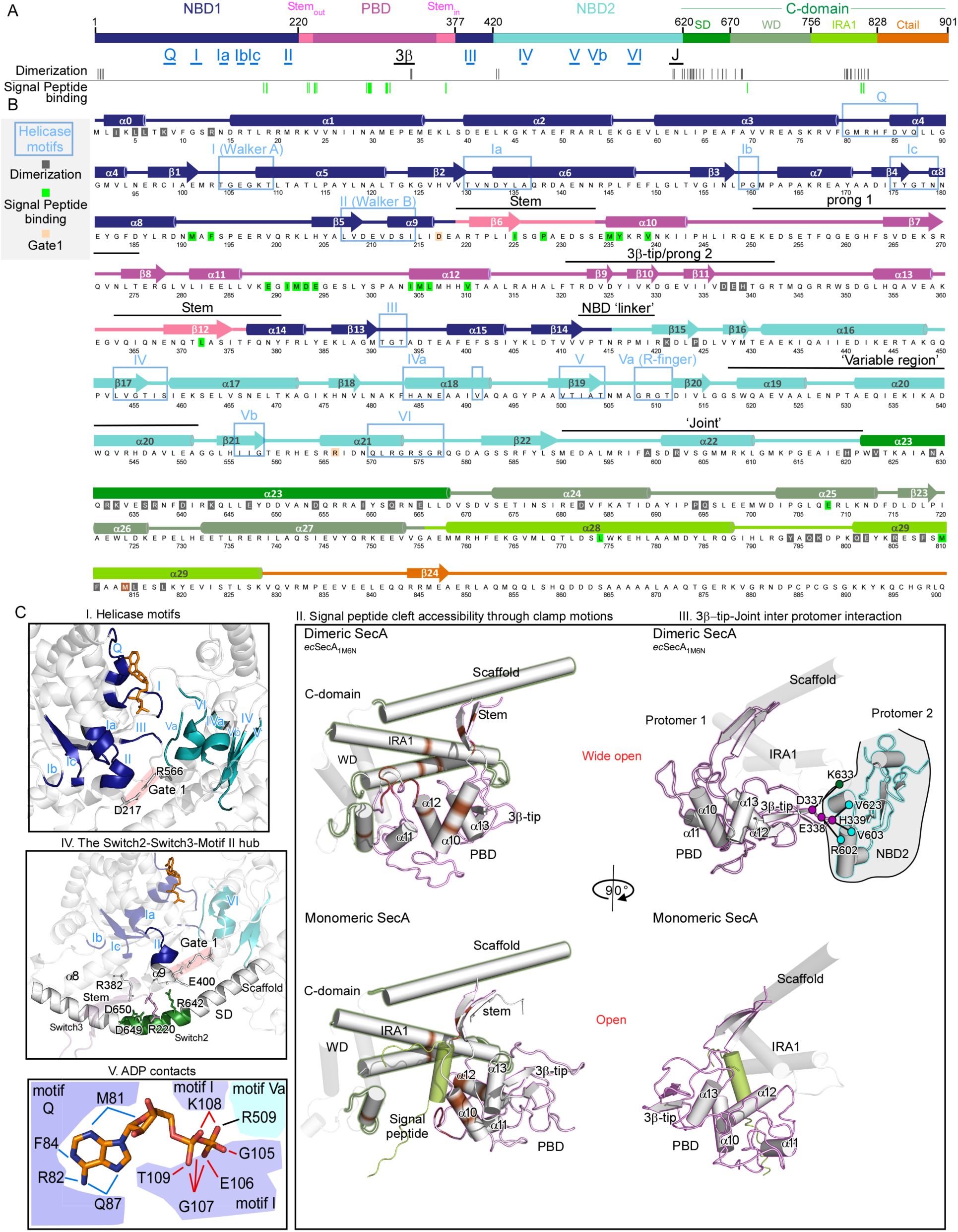
Map, sequence and structural elements of SecA (related to Introduction and Figures 1 and 2) **A.** Linear map of the domain organization of SecA. SecA consists of 4 domains: Nucleotide Binding Domain 1 (NBD1; dark blue); Preprotein Binding Domain (PBD; magenta) ‘sprouts’ out of NBD1 via two flexible linkers (Stem_out_ and Stem_in_; pink). Nucleotide Binding Domain 2 (NBD2; cyan), together with NBD1 form the ATPase motor. The C-domain consists of 4 subdomains; Scaffold Domain (SD; dark green), Wing Domain (WD; pale green), Intramolecular regulator of ATPase 1 (IRA1; lime green) and the flexible C-terminal tail (orange). Helicase motifs (in blue) and important structural elements (in black): 3β-tip and joint, are indicated. Dimerization residues were derived from dimeric *ec*SecA modelled after the *B.subtilis* dimeric SecA structure (PDB: 1M6N) and are detailed together with their paired residues in the supplementary Table S1 of (Gouridis et al., 2013). Residues that bind signal peptide were derived from (Gelis et al., 2007). **B.** Linear map of the primary sequence of SecA with secondary structural elements (coloured according to the domain organization in Panel A) and elements of functional or structural importance (adapted from (Papanikolau et al., 2007)). Blue rectangles: helicase motifs ((Papanikolau et al., 2007) and updated from (Fairman-Williams et al., 2010; Linder and Jankowsky, 2011)); cylinders: α-helices; arrows: β-strands. **C.** Blow ups of structural elements of SecA. **I.** Conserved Superfamily 2 helicase motifs (light blue rectangles in Panel B) (Linder and Jankowsky, 2011; Papanikolau et al., 2007) are mapped, onto the blow-up structure of the SecA nucleotide binding cleft. The open motor conformation of the ecSecA X-ray structure is shown (PDB ID: 2FSI). Helicase motifs in NBD1 and NBD2 are colored blue and cyan respectively (as in Fig. S1A), all other regions are in grey and rendered transparent. D217 and R566 (grey sticks) form the ‘Gate1’ salt bridge (shaded pink) that is required to break before the ATPase catalytic function can be activated (Karamanou et al., 2007). ADP is represented as orange sticks. **II.** Blow up image of the preprotein binding domain (PBD; outlined in magenta) and C-domain (outlined in green), with the clamp in the wide-open state (top panel; in dimeric SecA; *ec*SecA modelled after *bs*SecA; PDB ID: 1M6N) and open state (bottom panel; in monomeric SecA; PDB ID: 2VDA). SecA residues that interact with the signal peptide are coloured brown (Gelis et al., 2007). The signal peptide is shown in light green (bottom panel). Secondary structural elements of the PBD are indicated. **III.** Same as in panel II, with the images turned 90° clockwise. Top panel: In dimeric SecA (*ec*SecA modelled after *bs*SecA; PDB ID: 1M6N), 3β-tip residues (magenta circles) in the PBD, move under the IRA1 helices to make inter-protomeric contacts with the joint (cyan circles) and SD (green circles) of the second protomer. NBD2 of the second protomer is outlined in cyan. Bottom panel: Monomeric SecA (PDB ID: 2VDA) with the clamp in the open position shows the 3β-tip rotated away from IRA1 helices. **IV.** The Stem/Scaffold-Motif II Hub. The long α23 scaffold helix that connects NBD2 to the C-domain and regulates NBD function (PDB ID: 2VDA) contains a kinked middle (residues 640-650; green) that is a hotspot for inter-domain salt bridges. The Stem and Scaffold (coloured as in S1C.I and IV) contact each other through a hydrophobic core and salt bridges between D649-R220 and D650-R382. R642_Scaffold_ salt bridges with E400_NBD1_. Scaffold also participates in SecA dimerization (Fig. 2D; S1B) and becomes a prominent interaction site through which monomeric SecA binds SecY (Zimmer et al., 2008). **V.** SecA residues that contact ADP (dark grey backbone, CPK colouring) are labeled (PDB ID: 2FSI; from (Papanikolau et al., 2007)) and shown in connection to the helicase motif to which they belong. NBD1 residues are in blue, NBD2 residues are in cyan. Residues from motif Q mainly contact the adenosine (blue bonds) while residues from Motifs I (red bonds) and V (black bond) mainly contact the α and β phosphates.

**Figure S2.**
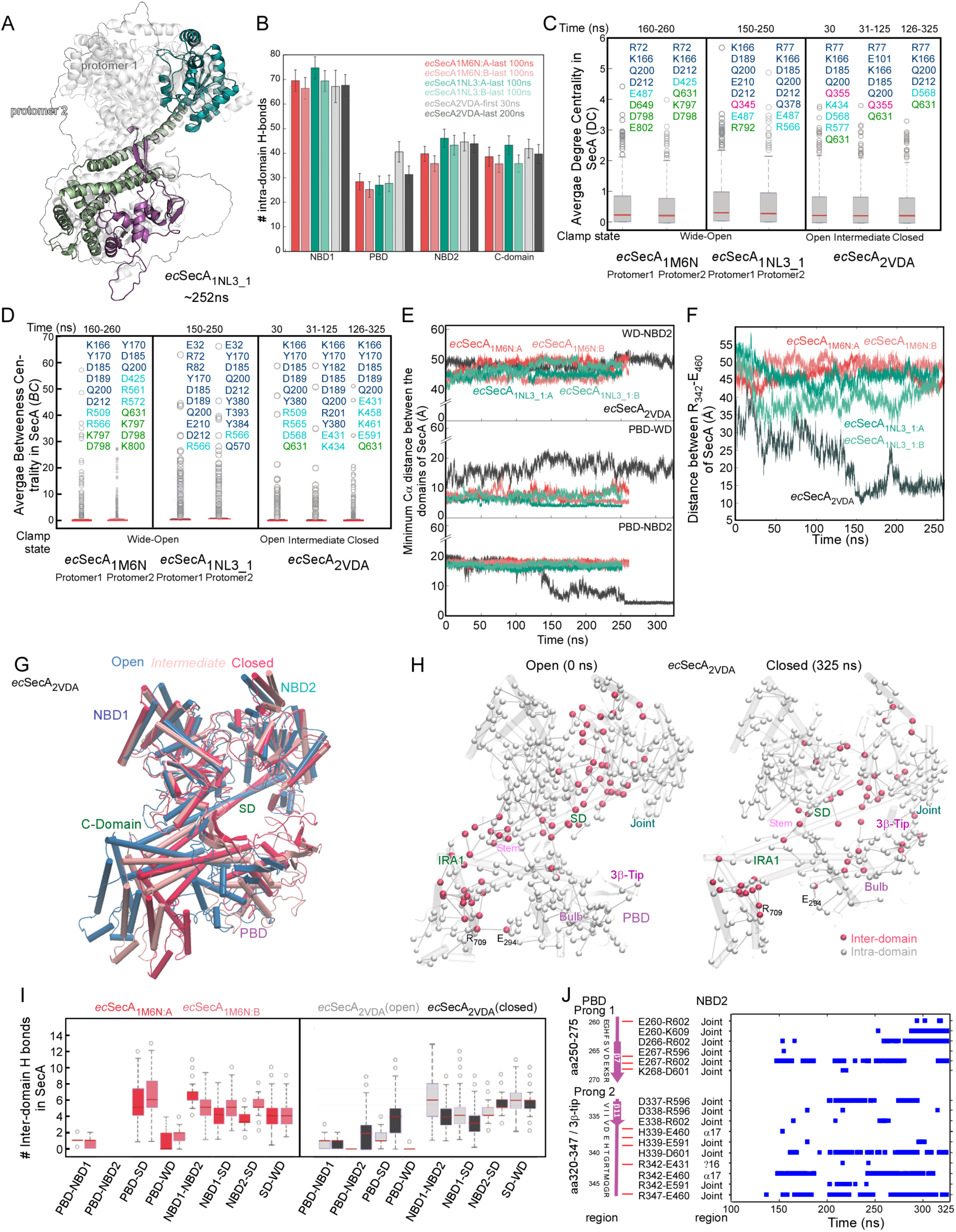
MD simulation of motions in dimeric and monomeric SecA and Graph analysis (related to Figures 1, 2 and 5). **A.** Two coordinate snapshots (0 and 252 ns) from the MD simulation for the dimeric *ec*SecA were visualized on the *M. tuberculosis* crystallographic dimer [PDB: 1NL3_1; (Sharma et al., 2003)] that was identified as being formed between adjacent asymmetric units in the crystal (Gouridis et al., 2013). The two snapshots were aligned based on the NBD1 region, using the MultiSeq plugin of VMD (Roberts et al., 2006), and superimposed. The front-facing protomer of time = 0 ns is gray, the one of time = 252 ns is colored (PBD: magenta; NBD2: cyan; C-domain: green; (Humphrey et al., 1996). In either case, the back facing protomer is presented only as an outline. Molecular graphics were prepared using Visual Molecular Dynamics, VMD (Humphrey et al., 1996). **B.** Average number of intra-domain H bonds for domains of indicated SecA states. In monomeric SecA (ecSecA2VDA) the first 30 ns (grey bar) and last 200 ns (black bar) of simulation time are presented. In the 2 forms of dimeric SecA (1M6N:red shades, 1NL3_1: green shades) the last 100 ns of simulation time is presented for each protomer (A and B) of the dimer. The largest difference in H-bonds was observed in the PBD of monomeric SecA between early(grey bars) and later simulation time (black bar). In the 2 forms of dimeric SecA, intra-domain H-bonds do not vary across simulation time, but there are subtle differences between the two protomers within a dimer. Histograms were generated using MATLAB R2017b. Error bars represent the standard deviation of each group. **C.** Distribution of average Degree Centrality values (*DC*) in simulations of monomeric and dimeric SecA (complete dataset in Table S2) (Freeman, 1979; Lazaratos et al., 2020). *DC* describes the number of direct H-bonds a residue can form with unique partners during the simulation so max value is ∼4-5. Average DC values were calculated for each residue by calculating its DC value at each coordinate set averaging across all coordinate sets used for the analysis. High average DC values for an amino acid residue mean that this group is in an environment in which it forms several H-bonds with different partners and are therefore indicative of stabilizing interactions. When located centrally in a H-bond cluster, groups with high DC commonly also have high Betweenness centrality (BC; See Panel C) values and are part of clusters which are stable. The residues with the highest average *DC* values (≥3) are indicated. To facilitate comparison, they are ordered according to primary sequence and colour-coded according to their domain (defined in Fig. S1A and B). Most of them map in the helicase motor. We visualize specifically the ones around the Scaffold/Stem/Motif II hub in Fig. S1C.VII. Most of these residues are highly conserved (Table S4). **D.** Distribution of average Betweenness Centrality (*BC)* values in simulations of monomeric and dimeric SecA (complete dataset in Table S2) (Freeman, 1977; Lazaratos et al., 2020). *BC* describes the importance of a residue in a H-bond network in terms of the fraction of shortest and continuous H-bonded pathways that pass through the residue and connect it with unique H-bond partners during the simulation. High *BC* values for an amino acid residue mean that it acts as a hub and controls multiple H-bond connections around it and at a distance. Such residues are commonly conserved and/or essential for function (Amitai et al., 2004; Lazaratos et al., 2020). Groups with the highest *BC* values (>10) are considered outliers. Top 10 residues with the highest average BC values are indicated and ordered according to primary sequence. Others include residues in NBD2 and the PBD. These groups are at the crossroads of H-bond paths that inter-connect numerous other protein groups. Most of these residues are highly conserved (Table S4). **E.** Changes in the minimum C*α* distance (y axis; Å) between WD-NBD2 (green), PBD-WD (red) and PBD-NBD2 (blue) regions of *ec*SecA during simulations of monomeric *ec*SecA_2VDA_ (x axis; ns). We calculate the distances of Cα atoms of unique residue pairs that belong to different protein domains and we keep for each time of the simulation, the minimum Cα distance of the residue pair which is closer to each other among others. Note that, as the simulation progresses, the distance between the PBD and NBD2 decreases. The corresponding analysis was carried out for the SecA dimers under the same conditions and force field is included for comparison (as indicated). Dimeric forms do not show the fast-conformational transition and re-orientation of the PBD seen in the monomer. This suggests that stability of the PBD in the Wide-open state in the same force field in the dimers is due to additional interactions that the dimers provide (Fig. S1C.V; Table S1; Table S2). **F.** Time series of Cα-Cα distances (y axis; Å) between R342_PBD_ and E460_NBD2_ of monomeric SecA (black line; *ec*SecA_2VDA_) or two structural variants of dimeric SecA (red and green lines; *ec*SecA_1M6N_ and *ec*SecA_1NL3_1_ respectively) during the MD simulation time (x axis; ns). For dimeric SecAs, distances were plotted for both protomers. These timeseries indicate that closing of the PBD in monomeric *ec*SecA associates with formation of a salt bridge between R342 and E460 (see Panel I below). In the two dimers, the salt bridge is not sampled, as the PBD remains away from NBD2 **G.** Open, Intermediate and Closed conformations (colour-coded as indicated) of the monomeric *ec*SecA_2VDA_ structure as illustrated by the MD simulation. Coordinate snapshots from the simulation were structurally aligned based on their NBD1 region and superimposed using the MultiSeq plugin of VMD (Roberts et al., 2006). **H.** Changes in inter- and intra-domain H-bonding interactions during a 325 ns MD simulation of *ec*SecA (PDB: 2VDA; 0 ns: Open state; 325 ns: Closed state). Cα atoms involved in inter-domain hydrogen bonds are depicted as red spheres, those involved in intra-domain bonds as white ones, against a grey cartoon backdrop. Lines between C*α* atoms represent H-bond distances. For simplicity, we only show unique H bonds between two protein groups. Here, the WD and SD sub-domains of the C-domain are considered as independent domains and “inter-domain” H-bond interactions between them are shown. **I.** Average number of inter-domain H-bonds (y axis) between the indicated domain pairs (x axis) during the MD simulation are represented as box plots (generated using MATLAB R2017b) for both monomeric and dimeric forms of SecA. For monomeric SecA, (ecSecA_2VDA_), H-bonds were calculated for the first 30 ns (grey bars) and last 200 ns (black bars) of the MD simulation. For dimeric SecA (ecSecA_1M6N_, ecSecA_1NL3_1_) H-bonds were calculated for the last 100 ns for both protomers (coloured as indicated). The central line of the box plot represents the median value. Bottom and top edges of the box indicate the 25th and 75th percentiles, respectively. An outlier (circles) is a value that is >1.5 times the interquartile range (1.5 of IQR) away from the top or bottom of the box. **J.** Inter-domain salt bridges (blue bars) between the indicated residues (y-axis) of *ec*SecA_2VDA_ were plotted as a function of the MD simulation time (x axis). Prong 1 (residues 250-275) and Prong 2 (residues 320-347) of PBD salt bridge with NBD2 residues of the second protomer during the course of the MD simulation. A salt bridge is considered to be formed if the distance between any of the oxygen atoms of acidic residues (ASP, GLU) and the nitrogen atoms of basic residues (ARG, HIS, LYS) are within 3.2 Å. Histidines in our simulations are singly protonated and thus neutral. We do not know if His339 can become doubly protonated and basic.

**Figure S3.**
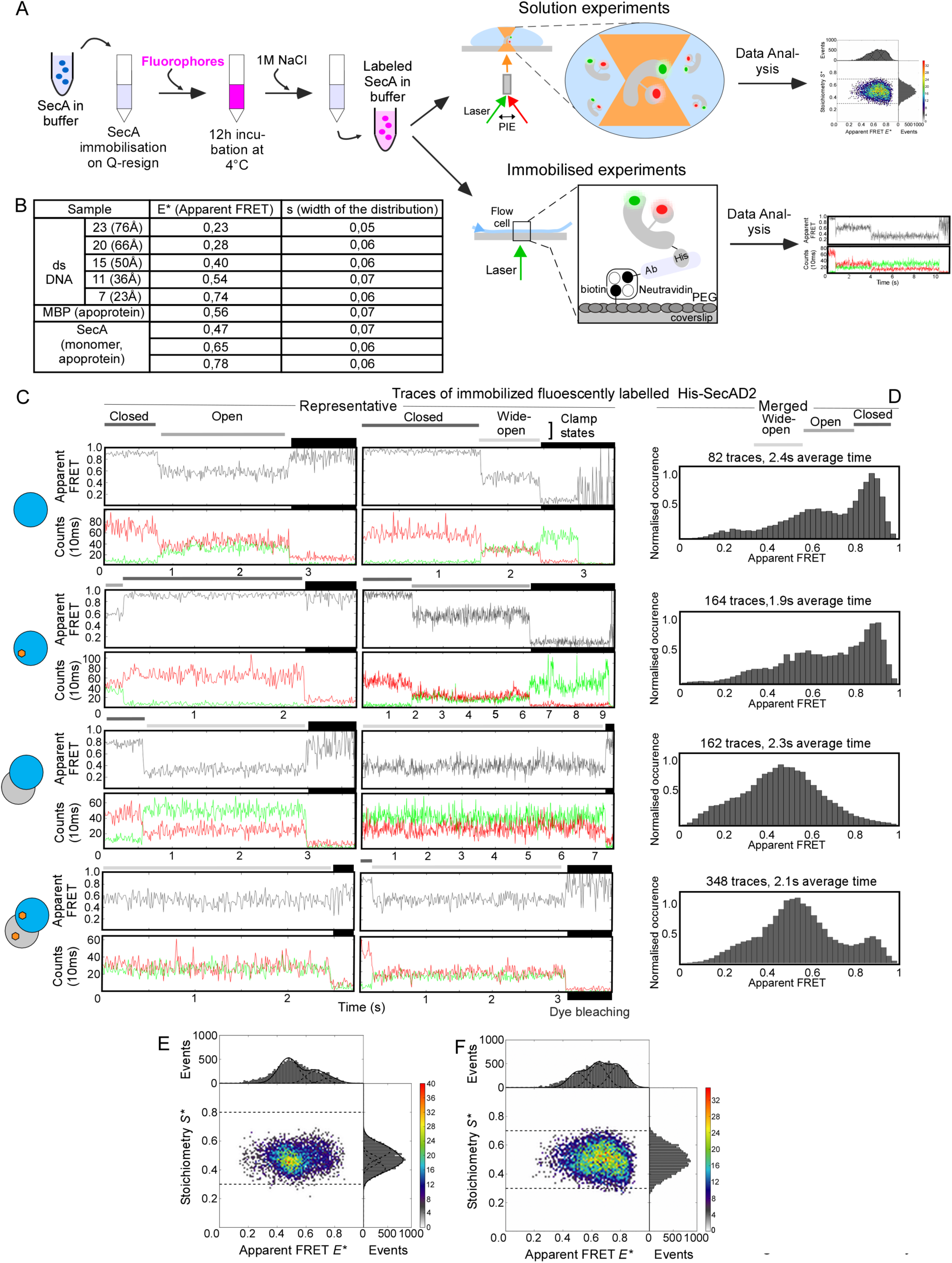
Single molecule FRET (smFRET) analysis in solution or with immobilized SecA molecules (related to Figures 1, 2, 3, 4 and 5). **A.** Schematic representation of smFRET workflow using SecA either in solution or immobilized. His-SecAD2 variant (V280C_PBD_; L464C_NBD2_) was stochastically labelled with Alexa555 (donor) and Alexa647 (acceptor) and used (50-100pM) to probe clamp dynamics. Solution experiments were carried out on a confocal pulsed interleaved excitation microscope. FRET values of SecA molecules randomly diffusing through the confocal volume were calculated and presented on 2D-histograms. For experiments using immobilized molecules, labeled SecA molecules were surface-immobilized with PEG-biotinylated-α-His antibody and their FRET trajectories were calculated and plotted vs time. **B.** Control FRET measurements using labelled DNA and protein standards. Doubled stranded (ds) DNAs were chemically synthesized with Alexa555 and Alexa647 fluorophores incorporated in positions that differed by 7 bp (23 Å), 11 (36 Å), 15 (50 Å), 20 (66 Å) or 23 (76 Å). smFRET measurements with these molecules were used to correlate Apparent FRET (E*) values with distance and distribution width (σ). Maltose Binding Protein (MBP) was purified and labeled (Alexa555, Alexa647) as described (de Boer et al., 2019). As MBP_Apo_ acquires one state, its FRET values were used to corroborate the dsDNA measurements. Extreme smFRET values have the smallest σ (=0.05) while medium FRET values have a bigger σ (=0.07). **C.-D.** Representative single traces (C), and projection of the indicated amount of single traces in the same graph (D), of smFRET experiments using immobilized His-SecAD2 molecules under the same conditions (indicated on the left pictogram). Grey-coded lines above graphs indicate different clamp states. The pictogram on the left indicates experimental conditions; single circle: monomer SecA; double circles: dimer SecA; blue: labelled protomer; grey: unlabeled protomer; orange: ADP. **C.**: Apparent FRET efficiencies (E*; y axis) for the fluorescence trajectories of HisSecA-D2 (black line) in the course of time (x axis). Donor (Alexa555; green) and acceptor (Alexa647; red) photon counts, recorded in parallel, were binned with 10 ms (y axis) and are shown in the course of time (x axis), below the Apparent FRET panels. **D.**: The number of single traces that are projected and their average time are indicated on top of each graph. **E.-F**. Pulse Interleaved Excitation (PIE)-based 2D plots derived from smFRET experiments of freely diffusing, monomeric (E) or dimeric (F) SecA. The x axis represents the apparent FRET value (E*), the y axis stoichiometry (S*) (see material and methods). Histograms, produced by plotting the apparent FRET values (E*) of multiple recordings (i.e. events), were fitted with three Gaussian distributions each having the width (σ) of the corresponding apparent FRET distribution (E*) (see B). Experiments using solution or immobilized molecules both agree that, the Wide-open state is enriched upon dimerization of SecA; variations of the plots might be due to lower statistics of the immobilized data.

**Figure S4.**
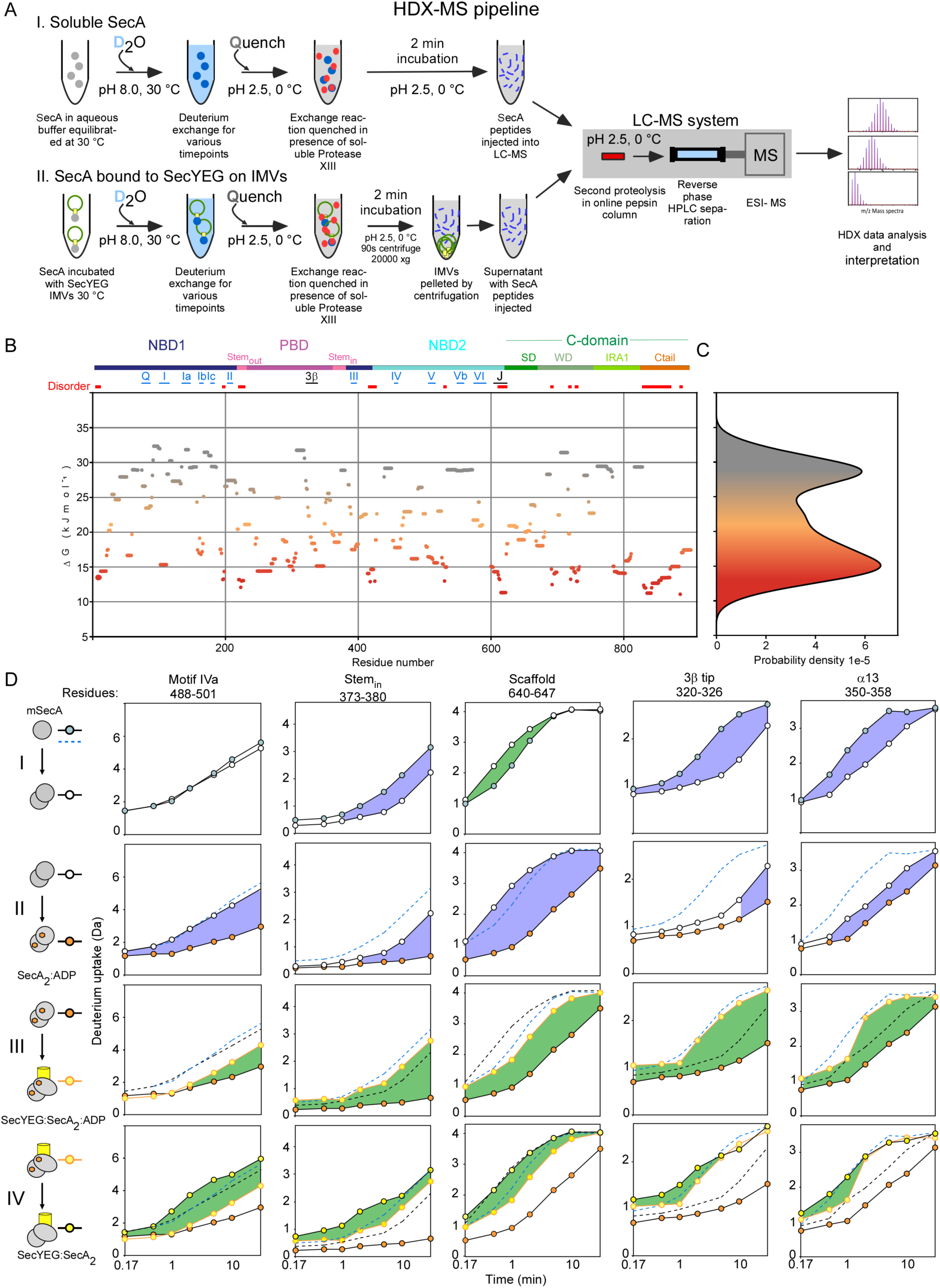
HDX-MS methodology, SecA Deuterium exchange rate map and selected D-exchange plots (related to Figures 1-7). **A.** Schematic representation of HDX-MS workflow using freely diffusing SecA (I) or bound to SecYEG-IMVs (II). I: Soluble SecA (grey circles) is equilibrated at 30°C. D-exchange (blue shading) is initiated by dilution in deuterated buffer F (1:10 dilution for 90% final D_2_O concentration) and incubated for various timepoints (10 s, 30 s, 1 m, 2 m, 5 m, 10 m, 30 m, 1440 m). The reaction is quenched with formic acid (0.7% V/V final concentration) at 4°C. Protease XIII (1 mg/mL) (grey shading) is added and the sample is incubated for 2 min on ice. II: In experiments with channel-bound SecA, soluble SecA (grey squares) is incubated with SecYEG-IMVs (green circles with yellow boxes) at a ratio of 1.5:1 (SecYEG:SecA_2_). An excess of SecYEG ensures all available SecA is bound to SecYEG. D-exchange is initiated, and the reaction is quenched as in solution experiments. During the quench step, at low pH, the SecYEG:SecA_2_ complex tends to dissociate (data not shown) and the free SecA is proteolyzed while SecYEG-IMVs are inaccessible to the protease. SecYEG-IMVs are pelleted by centrifugation, at 20,000 x g, (90s, 4°C), while SecA peptides (blue lines) remain in the supernatant. Sample handling between centrifugation and injection was maintained to a maximum of 30 s and was always at 4°C. I. and II.: The reaction is then injected into a LC-MS system, maintained at pH 2.5 and 0°C, and becomes further digested using a home-packed online pepsin column (Sigma-Aldrich) before peptides are separated by reverse phase HPLC. HDX-MS data were analyzed and interpreted as described in experimental methods. **B.** ΔG_ex_ values were calculated for each residue of SecA_2_ on PyHDX software (Smit et al., 2020) using time-course D-exchange experiments (see experimental methods), plotted from N-to C-terminus and coloured on a linear scale from grey (29 kJmol^-1^; rigid) to red (13 kJmol^-1^; disordered) as in Fig 1F. Flexible regions are in orange and fall in the middle of this range of ΔG_ex_ values (21 kJmol^-1^). Disordered islands (<13 kJmol^-1^) are indicated above the graph (red). The domain organization of SecA is indicated as a colored bar above the graph (colors as in Fig 1A). Lines depict the location and length of helicase motifs I-VI (blue), 3β-tip and the Joint (black) in the primary sequence of SecA. **C.** Histogram depicting the population density (x-axis) of residues as a function of ΔG_ex_ (y-axis aligned with S4B). The histogram is coloured based on ΔG_ex_ values as in B. **D.** HDX-MS kinetic plots compare the D-uptake of selected SecA peptides (as indicated on top of panels) in different states. I.-IV: Pictograms, on the left of panels, indicate the activation cascade of SecA as well as the current state pair comparison. Single circle: monomer SecA; double circles: dimer SecA; orange: ADP; yellow SecYEG. The symbols used on the kinetic plots for each state, are indicated on the right of the pictogram. Each row (I-IV) compares the D-uptake values (in Da) of the control state (top pictogram) to test state (bottom pictogram) across seven timepoints (10 s, 30 s, 1 m, 2 m, 5 m, 10 m and 30 m). In all plots, the y axis is scaled to equal the 100% deuteration control of each peptide (FDC, Table S1), the x axis is in logarithmic scale. Each point is the average of 3 replicate experiments. Standard deviation error bars (maximum of ±0.12 Da, average of ±0.06 Da) are within the area of the data point and hence were omitted. *Peptide identity:* Selected peptides, with the indicated residue numbers, span: Motif IVa (I, FDC= 7.97 Da), Stem_in_ (II, FDC= 4.79 Da), Scaffold (III, FDC= 4.22 Da), 3β-tip (IV, FDC= 2.63 Da) and α13 (V, FDC= 3.76 Da). *Colouring:* Purple: decreased, green: increased dynamics, in the test state of the indicated peptide compared to the control state. The greater the shaded area, the larger is the difference in dynamics between control and test states. I.: SecA_2_ (white circles, test) is compared to mSecA (blue circles, control). II.: SecA_2_:ADP (orange circles, test) is compared to SecA_2_ (white circles, control). ADP is maintained at 2 mM. mSecA data (blue dashed line) are included for comparison. III.: SecYEG:SecA_2_:ADP (yellow circles outlined orange, test) is compared to SecA_2_:ADP (orange circles, control). SecYEG:SecA_2_ is maintained at a molar ratio of 1.5:1. SecA_2_ data (black dashed line) are included for comparison. IV.: SecYEG:SecA_2_ (yellow circles, test) is compared to SecYEG:SecA_2_:ADP (yellow circles outlined orange, control).

**Figure S5.**
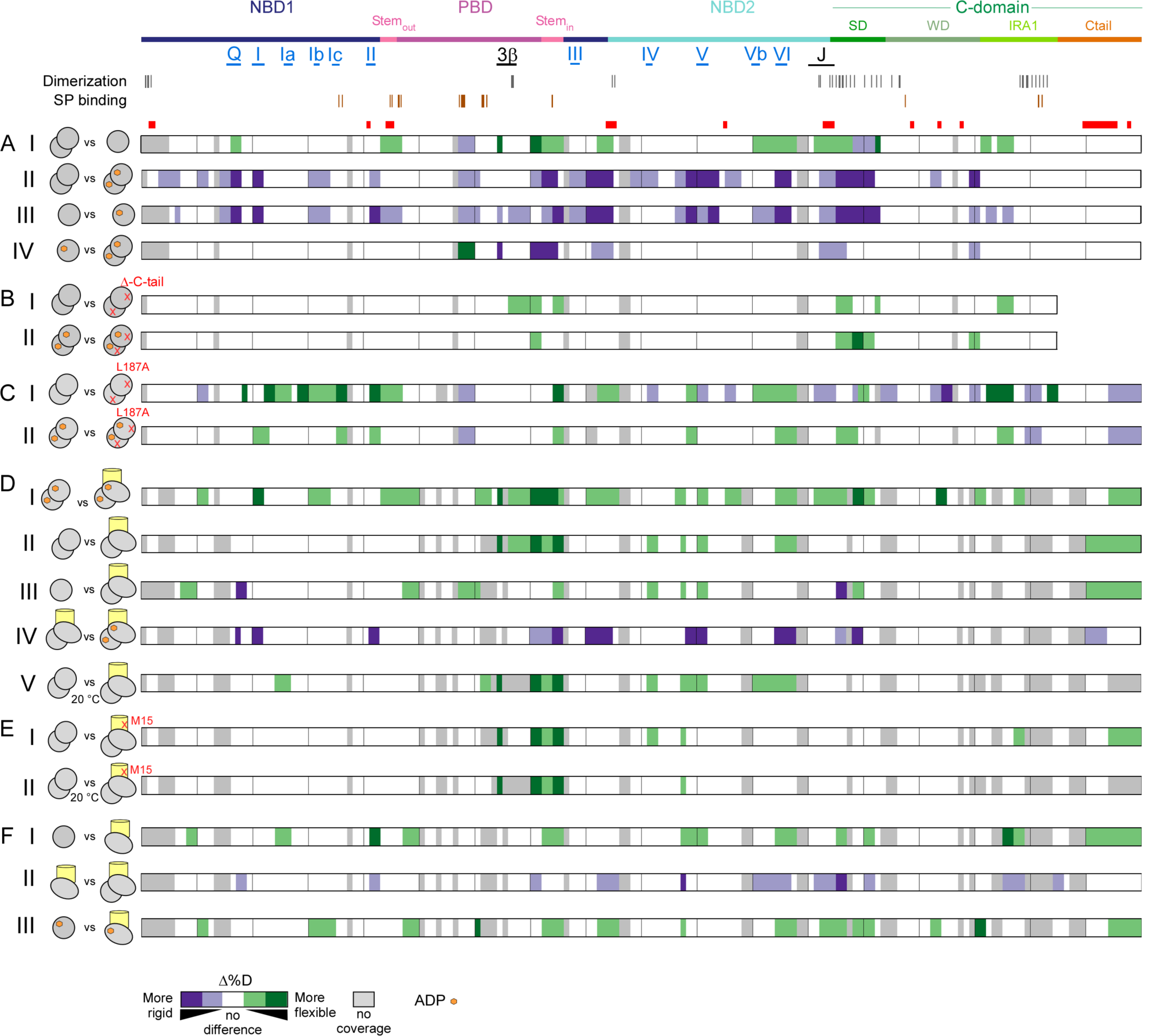
Δ%D comparison of localized dynamics in SecA and derivatives in various liganded states (related to Figures 2-6). **A.-H.** HDX-MS data from different conditions are compared pairwise as indicated on the pictograms (see below). Differences (**Δ)** in %D are colored (as explained below) on linear maps of SecA, from the N-to the C-terminus. *Above the Panels:* The domain organization, important structural elements (Helicase motifs; 3β-tip; Joint) and regions of disorder as well as their position, are indicated onto the linear map of SecA (as in Fig. S4B). *Left of Panels:* Pictograms indicate the current state pair comparison. Left pictogram: control; right pictogram: test state; grey single circle: monomer SecA; grey double circles: dimer SecA; orange hexagon: ADP; yellow cylinder: SecYEG; green cylinder: signal peptide; A red cross marks SecA or SecY mutants (as indicated). *Colouring of linear maps (Panels A-H):* Purple: decreased; green: increased dynamics of the ‘test state’ when compared to the ‘control state’. Colour shades indicate minor/major differences in dynamics (as depicted at the bottom of the Figure). All colored regions satisfy the significance threshold criteria (see Experimental methods). Grey: no coverage. **A.** Changes in the dynamics (Δ%D) of mSecA upon dimerization or/and ADP binding. I. SecA_2_ (control) is compared to mSecA (test). II. SecA_2_ (control) is compared to SecA_2_:ADP (test). III. mSecA (control) is compared to mSecA:ADP (test). IV. mSecA:ADP (control) is compared to SecA_2_:ADP (test). **B.** Changes in the dynamics (Δ%D) of SecA_2_ upon deletion of its C terminal tail, in the presence or absence of ADP. I. SecA_2_ (control) is compared to SecA_ΔCtail_ (test). II. SecA_2_:ADP (control) is compare to SecA_ΔCtail_:ADP (test). **C.** The effect of mutating Stem (i.e. L187A) on the dynamics (Δ%D) of SecA_2_ is examined, in the presence or absence of ADP. I. SecA_2_ (control) is compared to SecA(L187A) (test). II. SecA_2_:ADP (control) is compared to SecA(L187A):ADP (test). **D.** Changes in the dynamics (Δ%D) of SecA_2_ upon channel binding, in the presence or absence of ADP. I. SecA_2_ (control) is compared to SecYEG:SecA_2_ (test). II. mSecA (control) is compared to SecYEG:SecA_2_ (test). III. SecA_2_:ADP (control) is compared to SecYEG:SecA_2_:ADP (test). IV. SecYEG:SecA_2_ (control) is compared to SecYEG:SecA_2_:ADP (test). V. SecA_2_ (control; D exchange carried out at 20°C) is compared to SecYEG:SecA_2_ (test, D exchange carried out at 20°C). **E.** Changes in the dynamics (Δ%D) of SecA_2_ upon binding to SecY_M15_EG. I. SecA_2_ (control) is compared to SecY_M15_EG:SecA_2_ (test). II. SecA_2_(control; deuterium exchange carried out at 20°C) is compared to SecY_M15_EG:SecA_2_ (test, D exchange carried out at 20°C). **F.** Changes in the dynamics (Δ%D) of mSecA upon channel or/and signal peptide binding, in the presence or absence of ADP. I. mSecA (control) is compared to SecYEG:mSecA (test). II. SecYEG:mSecA (control) is compared to SecYEG:SecA_2_ (test). III. mSecA:ADP (control) is compared to SecYEG:mSecA:ADP (test).

**Figure S6.**
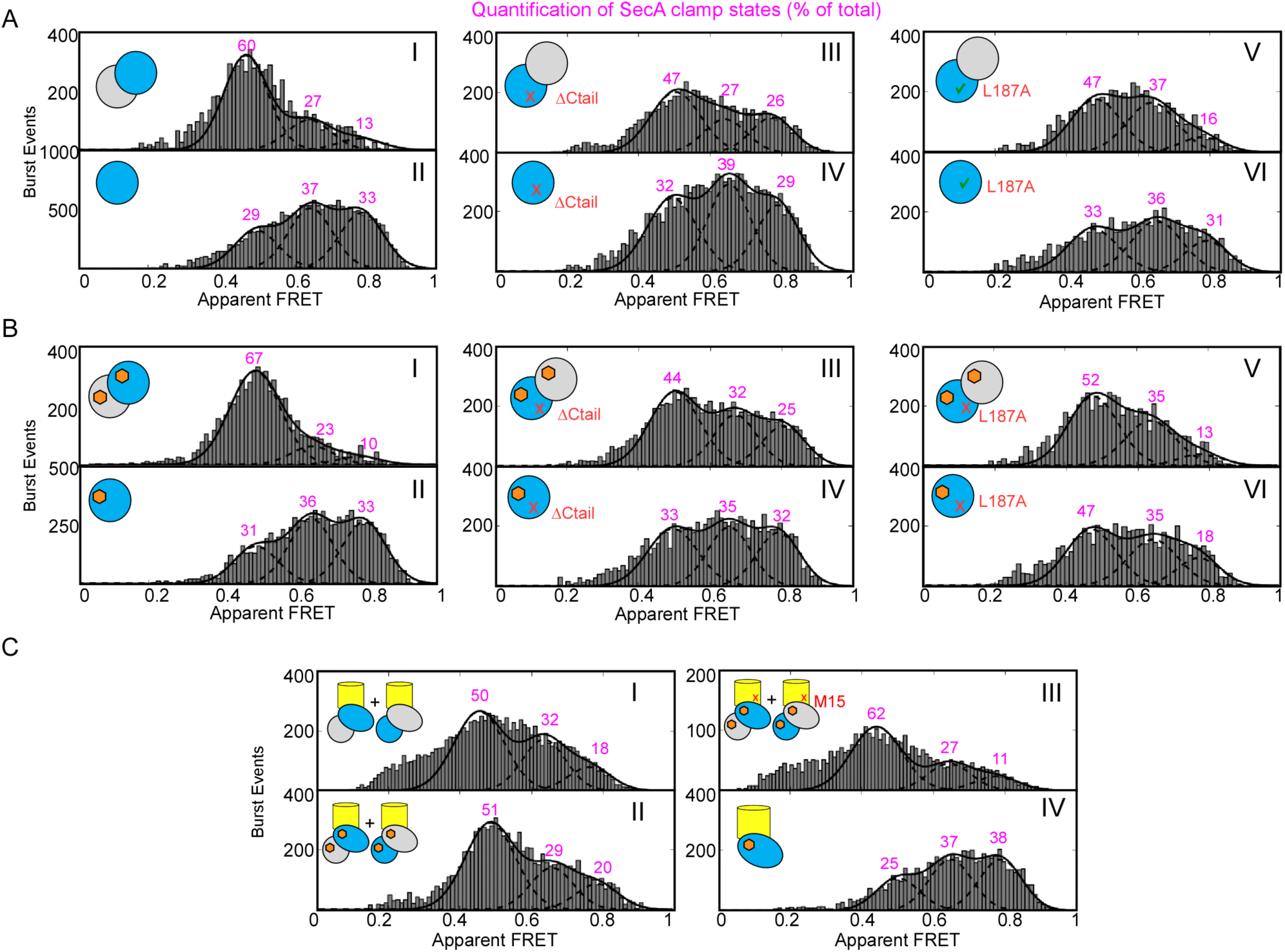
Intrinsic and ligand-modulated clamp dynamics in SecA determined by smFRET experiments in solution (related to Figures 1-6). **A.-C:** PIE-based 2D plots derived from smFRET experiments in solution, using SecA under the conditions indicated in the inlet pictograms (see below). Histograms [x axis: apparent FRET value (E*); y axis: burst events] were produced and fitted with Gaussian distributions as described in Fig. S3. The area under each curve was quantified and expressed as a percentage (pink numbers on top of curves) of the sum of areas under all curves. The labeled monomer of SecA (blue) was observed by using low pM concentration (50-100pM, kinetic monomer). For dimeric SecA, 500-1000nM non-labeled SecA (grey) was added to the labeled kinetic monomer and incubated for 30 min, 4°C. ADP was added (as indicated) at 1mM and incubated for 5 min, 4°C. For channel bound SecA, 1500 nM of SecYEG-IMVs were added to SecA (as indicated) and incubated for 30 min, 4°C. Signal peptide was added to free or channel bound SecA (as indicated), at 37µM and incubated for 5 min, 4°C. *Inlet pictograms indicate*: the fluorescent molecule(s) under observation (wild type or the indicated derivative), the state of SecA (monomer or dimer; apoprotein or ADP bound; freely diffusing or channel bound), the monodispersity of sample (for freely diffusing SecA molecules) or the ensemble of molecules (for channel bound SecA molecules), SecA homodimers (wild type) vs heterodimers (the labelled SecA protomer is the indicated mutant while the non-labelled protomer is wild type), the activated SecA protomer upon channel binding (oval) vs non-active protomer (circle), SecA or SecYEG mutants (red letters/cross). *Colors/shapes indicate:* Blue: labelled SecA protomer; grey: non-labelled SecA protomer; orange hexagon: ADP; yellow cylinder: SecYEG; green cylinder: signal peptide; circle: non-active channel bound SecA protomer; ovals: activated channel bound SecA protomer; red cross: protomer with mutation; pink: quantification (%) of SecA’s clamp states **A:**Freely diffusing SecA and mutant derivatives in apoprotein form; dimer SecA (I), monomer SecA (II), dimer SecA_ΔCtail_ (III), monomer SecA_ΔCtail_, dimer SecA(L187A), monomer SecA(L187A). **B:** Freely diffusing SecA and mutant derivatives in ADP bound form, shown as in A. **C:** Channel bound SecA; Channel-bound dimer SecA apoprotein (I), channel-bound ADP-bound dimer SecA (II), SecY_M15_EG bound ADP bound dimer SecA (III), channel-bound ADP-bound monomer SecA (IV)

**Figure S7.**
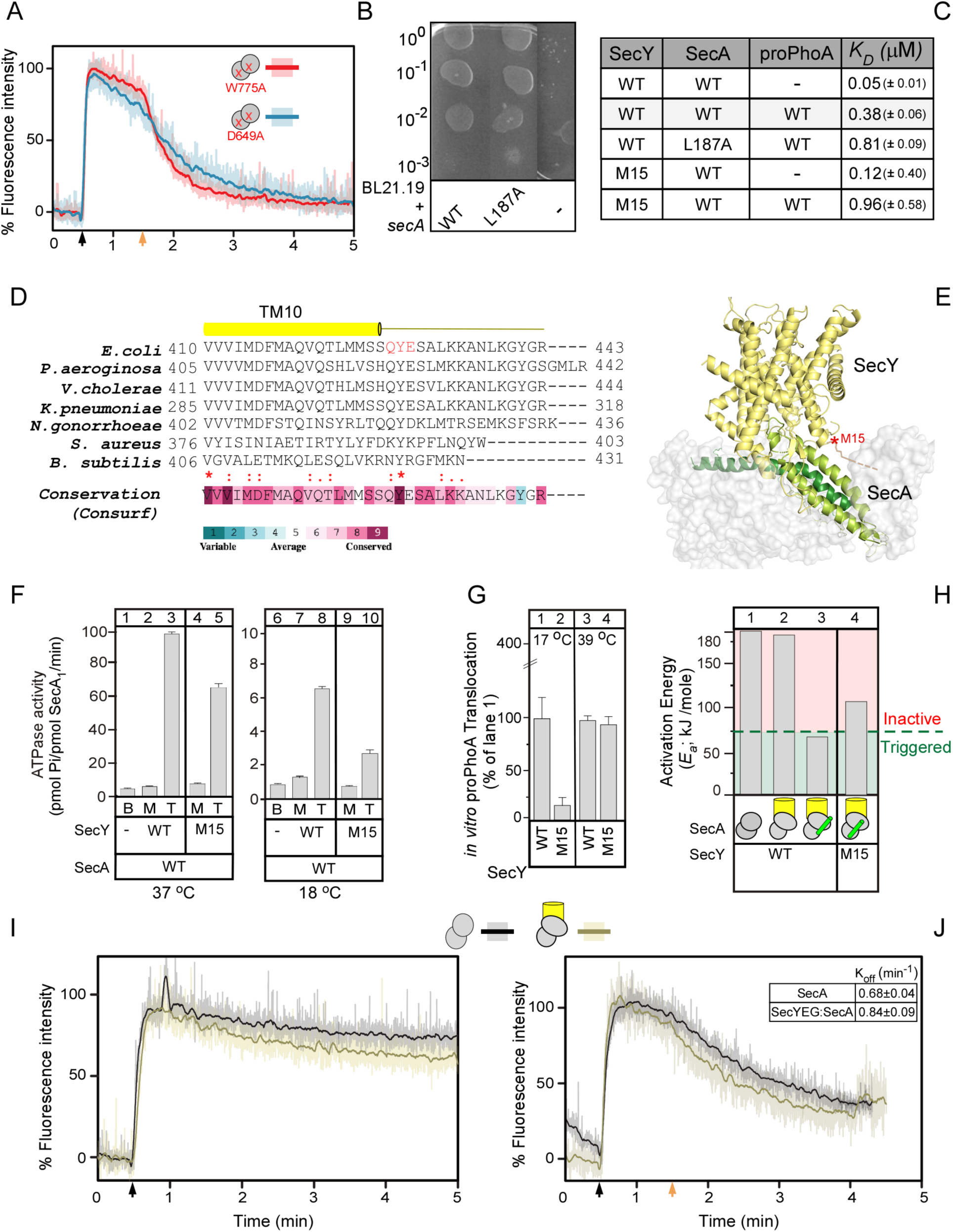
*In vitro* and *in vivo* functional properties of SecA and SecYEG derivatives (related to Figures 3-6). **A.** Binding of MANT-ADP to elevated ATPase mutants of SecA was monitored by fluorescence spectrometry. The low fluorescence of MANT-ADP in H_2_O increases upon binding to the hydrophobic environment of the nucleotide binding cleft of SecA (Galletto et al., 2000)(see Fig. 3E). The data is normalized taking the fluorescence signal of free MANT-ADP as 0% and that of SecA bound MANT-ADP as 100% (maximum fluorescence intensity). We monitored two previously described elevated ATPase mutants, SecA(D649A) (blue line) (Keramisanou et al., 2006) and SecA(W775A) (red line) (Vrontou et al., 2004). Raw fluorescence data (transparent lines) is superimposed with smoothened data (solid lines). 1 µM of SecA mutant was added to 1 µM of MANT-ADP at t= 30 sec (black arrow) and the sample was chased with 2 mM cold ADP at t= 90 sec. Curves represent the mean of three replicate measurements**B.** *In vivo* genetic complementation of the *E.coli* BL21.19 thermo-sensitive mutant strain by either an empty vector or one carrying the *secA* wt or mutants, as indicated. Serial dilutions of a culture (OD_600_=0.5) were spotted (12μl) on LB-Ampicillin plates and grown at 42°C. *n*= 3 biological replicates; a representative picture is shown. **C.** Binding affinities (K*_D_*) of SecA and derivatives for the SecYEG channel (as indicated; white rows) and of channel-bound SecA and derivatives for proPhoA (as indicated; grey rows). *n=* 6 biological repetitions; mean values (± SEM) are shown. SecA binds to SecY_M15_EG with high affinity and subsequently acquires a high affinity for preproteins, a demonstration that its association to SecY_M15_ rendered its two protomers functionally asymmetric. **D.** The C-termini of SecY from selected bacteria are aligned. TM10, the last transmembrane helix of SecY is indicated. The consensus sequence (bottom) was derived from alignment of 58 sequences from representative phyla across the domains of life (Karamanou et al., 2008) and analysis in the Consurf server of 300 homologues. Colour code for residue conservation: Red= 100%; Light blue= >70% identical or of similar property. += basic; @= acidic; s=small (A, G); h=hydrophobic (ILVMA). All residue numbering corresponds to the *E.coli* protein. Y429, a key residue, mutated in either the M15 derivative used here (Q_428_Y_429_E_430_-AAA)(Karamanou et al., 2008) or in SecY(Y429D) (Matsumoto et al., 2000; Matsumoto et al., 1997; Taura et al., 1997), renders SecY cryo-sensitive and the cells non-viable below 20°C. **E.** Structural model of the *E.coli* SecYEG:SecA complex generated by homology modelling against the SecYEG:SecA structure from *T. maritima* (PDB ID: 3DIN) (Vandenberk et al., 2019) reaching up to residue 427. Q_428_Y_429_E_430_, the site mutated on the M15 derivative (red asterisk) and the other last carboxy-terminal residues of SecY (yellow/grey drawn line), are not contained in the coordinates of the solved structures. These residues are likely to extend into making contacts with IRA1 (light green) and the scaffold domain (dark green). The last 13 C-terminal residues of *E.coli* SecYEG were shown to bind to SecA, using immobilized peptide arrays (Karamanou et al., 2008). **F.** The ATPase activity of the freely diffusing (basal; B; 0.4μM SecA), SecYEG channel-bound (membrane; M; 1 μM SecY) and translocating (T; SecY plus 9 μM proPhoACys^-^) SecA was determined as described (Gouridis et al., 2010). Mutant derivatives and incubation temperatures are indicated. *n=* 6 biological repeats; mean values (± SEM) are shown. **G.** *In vitro* translocation of proPhoACys^-^ (9μM) driven by wild-type or mutant translocases (i.e. wt SecA and the indicated SecY mutations), as described (Gouridis et al., 2010). The translocated material by the wild-type translocase at 17°C, or 39°C, is considered 100% for the indicated temperature. All other values were expressed as a percentage of the corresponding value. *n=* 3 biological repetitions; mean values (± SEM) are shown. **H.** The Activation Energies (*E_a_*) of wild type, or the indicated mutant, translocases under translocation conditions (0.4μM SecA; 1 μM SecYEG supplemented with 9 μM proPhoACys^-^, indicated by green tube) derived from Arrhenius plots, as described (Gouridis et al., 2009). *n=3* biological repeats. SecY_M15_ is cryosensitive (Karamanou et al., 2008); With the exception of the cryosensitive SecY_M15_ that exhibited an additional linear period from 20-28°C, all Arrhenius plots had two linear parts [0-20°C and 20-40°C; as described (Gouridis et al., 2009)]. **I-J.** Binding of MANT-ADP to SecA_2_ assembled in the translocase holoenzyme was monitored by fluorescence spectrometry. The data is normalized taking the fluorescence signal of free MANT-ADP as 0% and that of SecA_2_ bound MANT-ADP as 100% (maximum fluorescence intensity). Raw fluorescence data (transparent lines) is superimposed with smoothened data (solid lines). 1 µM of SecA_2_ or SecYEG:SecA_2_ (1.2:1 µM) were added at t= 30 sec (black arrow). In chase experiments (**J**), 2 mM cold ADP was added at t= 90 sec (orange arrow). Curves represent the mean of three replicate measurements. **I.** Fluorescence experiments comparing binding of SecA_2_ (black line) or SecYEG:SecA_2_ (brown line) to MANT-ADP. Fluorescence intensity was monitored for 5 min. **J.** Fluorescence experiments comparing binding of SecA_2_ (black line) or SecYEG:SecA_2_ (brown line) to MANT-ADP with a subsequent chase of 2 mM cold ADP (orange arrow). Fluorescence intensity was monitored for 4.5 mins. The off rate of MANT-ADP upon addition of cold ADP chase to both SecA_2_ and SecYEG:SecA_2_ was calculated using a one phase decay equation (Graphpad Prism).

### Supplemental tables

**Table S1** (related to Introduction and Figures 1, 2 and S1)

Analysis of SecA structures from the Protein data bank (in **XLS** file).

**I. X-ray structures information:** Summary information on X-ray structures of SecA homologues used in this study (organism, ligands, other factors, crystallographic conditions, the oligomeric state of the protein as identified in the crystal and the one that is proposed by EPPIC to be physiologically relevant (http://www.eppic-web.org/ewui/).

**II. All the PBD interactions:** All of the interdomain PBD interactions identified in different X-ray structures. Initially, homology models of *E. coli* SecA were created based on the available X-ray structures, using the Swiss-model web tool (https://swissmodel.expasy.org). Then, for every structure, all the interactions were calculated using the PIC webserver (http://pic.mbu.iisc.ernet.in) and only the interdomain PBD interactions were reported. In dimeric state of the protein, inter-protomeric interactions are also reported. The color of the interaction also indicates the nature of this interaction. Green: lipophilic; blue: hydrogen bond; red: ionic; purple: cationic-pi interaction. This table is further organized based on the oligomeric and the clamp state of the protein in four main groups (Dimer Wo, Monomer WO, Monomer O and Monomer C).

**III. Merged PBD interactions:** The table contains the merged interactions from different structures belonging at the same of the above-mentioned group. An extra column indicates the domain (color coded) from the respective amino acid residues.

**IV. Scaffold interactions:** The table contains all the interdomain scaffold interactions. In dimeric state of the protein, inter-protomeric interactions are also reported.

**Table S2** (related to Figures 1, 2, 5, S1 and S2)

Compiled data from atomistic MD simulations (in XLS file)

**I. Intra-domain H-bond**: H-bond frequencies (in %) between Intra-domain residue pairs for all MD simulations. Data are shown for 3 simulation time blocks of monomeric SecA (*ec*SecA_2VDA_) and the last 100 ns of simulation time for dimeric SecAs (*ec*SecA_1M6N_ and *ec*SecA_1NL3_1_). H-bond frequency, or occupancy, is defined as the percentage of length of the analyzed trajectory segment during which two residues are H-bonded. Data are shown for both protomers of dimeric SecA. Values above 20% are highlighted in red.

**II. Inter-domain H-bond**: H-bond frequencies (in %) between Inter-domain residue pairs during 325 ns simulation of monomeric SecA (ecSecA_2VDA_) and during 252-262 ns simulation of dimeric SecA in two different conformer (1. *ec*SecA_1M6N_, 2. *ec*SecA_1NL3_1_). For monomeric SecA H-bonding data is collated for the following simulation time blocks 1) first 30 ns, 2) 95 ns simulation time, 3) last 200 ns simulation time. For dimeric SecA H-bonding data is collated for the last 100 ns simulation time. H-bond frequency, or occupancy, is defined as the percentage of length of the analyzed trajectory segment during which two residues are H-bonded. Values above 20% are highlighted in red.

**III. Average DC**: Average Degree Centrality (DC) for all residues of SecA. Data is shown for both monomeric SecA (*ec*SecA_2VDA_) for 3 simulation time blocks and last 100 ns simulation time of dimer SecA (*ec*SecA_1M6N_ and *ec*SecA_1NL3_1_). Degree centrality (DC) of a node n gives the number of edges (H-bonds) connecting to n. The DC value of node n can be normalized by dividing its DC by the maximum possible edges to n (which is N-1, where N is the number of total nodes in the graph). In our analysis, we consider as nodes the amino-acid residues, edges the H bonds and paths the continuous chain of H-bonded residues. When a path between two nodes is the shortest, it has the least number of intermediate nodes (Lazaratos et al., 2020). Higher values indicate a higher propensity for forming stable H-bonds. Values above 2.5 are highlighted in red.

**IV. Average BC**: Average Betweenness Centrality (BC) for all residues of SecA. Data is shown for both monomeric SecA (*ec*SecA_2VDA_) for 3 simulation time blocks and last 100 ns simulation time of dimer SecA (*ec*SecA_1M6N_ and *ec*SecA_1NL3_1_. Betweenness centrality of a node n is given by the number of shortest paths that link any other two nodes (n1, n2) and pass via node n, divided by the total number of shortest paths linking n1 and n2. BC of node n can be normalized by dividing its BC by the number of pairs of nodes in the graph not including n. In our analysis, we consider as nodes the amino-acid residues, edges the H bonds and paths the continuous chain of H-bonded residues. When a path between two nodes is the shortest, it has the least number of intermediate nodes. High BC values for an amino acid residue mean that it acts as a hub and controls multiple H-bond networks around it and at a distance (Lazaratos et al., 2020). Values above 10 are highlighted in red.

**V. Average H-bond**: Average number of intra and inter domain H-bonds in indicated simulation time blocks for all MD simulations. Intra domain H-bonds are presented for all 4 domains of SecA. Inter domain H-bonds are presented for the indicated domain pairs. Data are shown for two types of calculations, H-bonds mediated entirely by residue-residue contacts (’protein’) and H-bonds that can also involve 1 water molecule to bridge two residues (1-water bridge).

**VI. DC (non-averaged)**: Degree centrality (DC) values for all residues of SecA. Data is shown for both monomeric SecA (*ec*SecA_2VDA_) for 3 simulation time blocks and last 100 ns simulation time of dimeric SecA (*ec*SecA_1M6N_ and *ec*SecA_1NL3_1_). Absolute DC values show number of H-bonds are residue is involved in during the indicated simulation time. Values above 10 are highlighted in red.

**VII. 1-w Intra-domain H bond:** H-bond frequencies (in %) between one-water bridged Intra-domain residue pairs for all MD simulations. One-water bridge is considered an H bond between two residue pairs mediated by one water molecule. Data are shown for 3 simulation time blocks of monomeric SecA (*ec*SecA_2VDA_) and the last 100 ns of simulation time for dimeric SecAs (*ec*SecA_1M6N_ and *ec*SecA_1NL3_1_). H-bond frequency, or occupancy, is defined as the percentage of length of the analyzed trajectory segment during which two residues are H-bonded. Data are shown for both protomers of dimeric SecA. Values above 20% are highlighted in red.

**VIII. 1-w Inter-domain H bonds:** H-bond frequencies (in %) between one-water bridged Inter-domain residue pairs during 325 ns simulation of monomeric SecA (*ec*SecA_2VDA_) and during 252-262 ns simulation of dimeric SecA in two different conformer (1.*ec*SecA_1M6N_, 2. *ec*SecA_1NL3_1_). One-water bridge is considered an H bond between two residue pairs mediated by one water molecule. For monomeric SecA H-bonding data is collated for the following simulation time blocks 1) first 30 ns, 2) 95 ns simulation time, 3) last 200 ns simulation time. For dimeric SecA H-bonding data is collated for the last 100 ns simulation time. H-bond frequency, or occupancy, is defined as the percentage of length of the analyzed trajectory segment during which two residues are H-bonded. Values above 20% are highlighted in red.

**Table S3** (related to Figures 1-7, S4 and S5)

HDX-MS Data table (in **XLS** file).

(Presented as recommended in (Masson et al., 2019). Data table contains raw D-uptake values for all states, percent uptake values and pairwise compared ΔD-uptake values between different experimental states)

**I. HDX summary Table**: Experimental details of all HDXMS results provided as recommended in Masson et al. 2019. Coverage map of SecA peptides obtained from solution (Fig S4A.I) and SecYEG bound SecA on IMVs (Fig. S4A.II) experiments.

**II. SecA all states raw data:** Raw D-uptake values (in Da) are shown with standard deviation values for all peptide in various states of SecA presented in this study. Peptide sequence and residue numbers are also shown. D-uptake values are shown as % D-uptake of the 100% deuteration control. % uptake values are coloured based on the provided colour key.

**III to XI**

Pairwise differential D-uptake values comparing various states of SecA are shown. Differential D-uptake between 2 states is shown both as raw ΔD (in Da) differences and %ΔD (difference in % uptake) values in separate columns. When comparing 2 states, the control state is in black and the test state is in red (always second) in the column header. ΔD differences greater than 0.5 Da are considered significant and coloured based on either protection (negative values, purple) or increased flexibility (positive values, green). A second threshold is applied based on Δ%D values; differences above 10% are considered significant. Peptides are considered significantly altered between 2 states if they satisfy both the ΔD and Δ%D thresholds (see experimental methods). Differences between 10%-20% are considered minor and differences greater than 20% are considered major differences. Protection is shown in shades of purple and increased flexibility is shown in shades of green (according to the provided colour key).

**III. SecA-Ligand:** Dimeric SecA in solution is compared to ADP-bound SecA.

**IV. mSecA-Ligand**: Monomeric SecA (mSecA) in solution is compared to dimeric SecA and ligand bound mSecA states:mSecA:ADP, SecA, SecA:ADP

**V. SecA-SecYEG-Ligands**: Dimeric SecA bound to SecYEG (SecYEG:SecA) is compared to the following states: SecA, SecA:ADP, mSecA, mSecA:ADP, SecYEG:SecA:ADP, SecYEG(M15):SecA.

**VI. SecA-SecYEG-SecYEG(M15)-18C**: Dimeric SecA in solution is compared to SecYEG:SecA and SecYEG(M15):SecA. Deuterium exchange for these experiments were carried out at 18°C

**VII. mSecA-SecYEG-Ligands**: Monomeric SecA (mSecA) bound to SecYEG (SecYEG:mSecA) is compared to the following states: mSecA, mSecA:ADP, SecYEG:mSecA:ADP, SecYEG:SecA, SecYEG:SecA:ADP

**VIII. SecAL187A**: SecA(L187A) in solution is compared to the following states: SecA(L187A):ADP, SecA, SecA:ADP

**IX. SecAdelCtail**: SecAΔCtail in solution is compared to the following states: SecAΔCtail:ADP, SecA, SecA:ADP

**Table S4** (related to Figure 1)

Amino acid conservation scores for all residues of *ec*SecA_2VDA_ from Consurf database (Ben Chorin et al., 2020). (in **DOC** file). Table contains conservation scores and residue variety for each residue of *E. coli* SecA. Table was generated by Consurf database using *ec*SecA_2VDA_ as the input structure.

### Supplementary Movies

The movies derived from crystallography and MD simulated states. Both movies were created as visualization models of the PBD motion. The movies were produced by using Pymol (https://pymol.org/) and composed in iMovie software.

**Movie S1** (related to Introduction and Figures 1 and S2)

For Movie S1, structural frames from the MD simulations (0 and 325ns) of monomeric SecA_2VDA_ were used.

**Movie S2** (related to Introduction, Figures 1, 2 and S2)

For Movie S2, the *E.coli* homology models (Sardis and Economou, 2010) of wide open, open and closed PBD states were used.

### Materials and Methods

#### List of Buffers

| Buffer | Composition |
| --- | --- |
| A | 50 mM Tris-HCl, pH 8.0, 1M NaCl |
| B | 50 mM Tris-HCl, pH 8.0, 50 mM NaCl |
| C | 50 mM Tris-HCl, pH 8.0, 50 mM NaCl, 50% v/v glycerol |
| D | 50 mM Tris-HCl, pH 8.0, 50 mM NaCl, 6 m Urea, 50% v/v glycerol |
| E | 50 mM Tris-HCl pH 8.0, 50 mM KCl, 1 mM MgCl<sub>2</sub>, 1 mM DTT |
| F | 50 mM Tris-HCl pH 8.0, 50 mM KCl, 1 mM MgCl<sub>2</sub>, 4 μM ZnSO<sub>4</sub>, 2 mM TCEP |
| G | 1.3% formic acid, 4 mM TCEP, 1 mg/mL fungal protease XII |
| H | 50 mM Tris-HCl pH 8.0, 50 mM NaCl, 0.1mM EDTA |
| I | 50 mM Tris-HCl pH 8.0, 1 M NaCl, 0.1mM EDTA |
| J | 50 mM Tris-HCl pH 8.0, 50 mM NaCl, 0.01mM EDTA |
| K | 50 mM Tris-HCl pH 8.0, 50 mM KCl, 1 mM MgCl<sub>2</sub> |

#### List of Strains

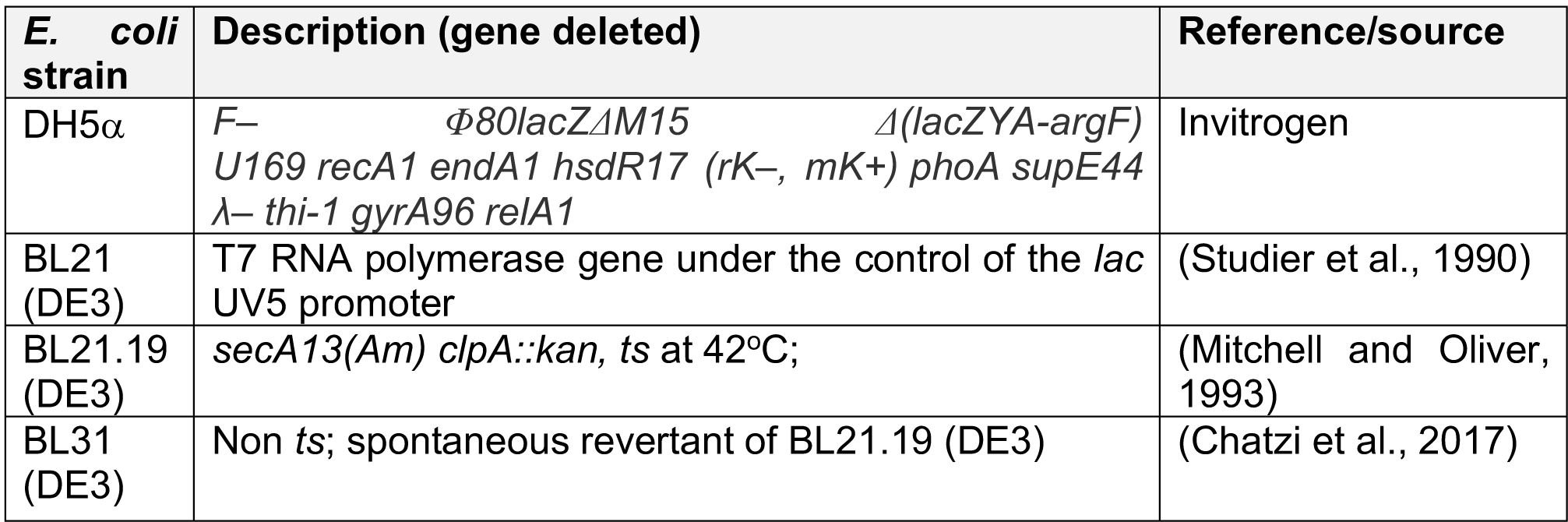

#### List of Plasmids

| Gene | Uniprot accessi on number | Plasmid name | Vector | Description/source/reference |
| --- | --- | --- | --- | --- |
| *secA* | P10408 | pIMBB1280 | pET3a | (Gouridis et al., 2013) |
| - |  | pLMB0081 | pET3a | pIMBB1280 was digested with NcoI, the *secA* N31-898 fragment was removed, the plasmid was re-ligated and used for *secA* NcoI fragment cloning |
| *secA*($\Delta\alpha 0/\alpha 1$-6A) | P10408 | pIMBB1286 | pET3a | or mSecA (Gouridis et al., 2013) |
| *secA*(V280C/L464) | P10408 | pLMB1646 | pET3a | or SecA-D2 (Vandenberk et al., 2019) |
| His *secA* cys- | P10408 | pLMB1791 | pET16b | *secA* N1-901 cys- (C98S/C885S/C887C/C896C) fragment from pLMB0092 [pET3A *secA* cys- (1-901)] was inserted in pET16b after NdeI-BamHI digestion |
| His *secA* cys- (V280C/L464) | P10408 | pLMB1819 | pET16b | or His-SecA-D2<br>The V280C/L464C mutations were introduced in pET16b His *secA* cys- (pLMB1791) using primer pairs X1824-X1825 and X1768-X1769 |
| secA N1-834 | P10408 | pIMBB1296 | pET5a | (Gelis et al., 2007) |
| His secA cys-N1-834 (V280C/L464C) | P10408 | pLMB2024 | pET16b | Two stop codons were introduced in pLMB1819 at position 835 using primer pairs X1024-X1025 |
| His secYEG | P0AGA2<br>P0AG96<br>P0AG99 | pIMBB336 | pET610 | Gift from A. Driessen, University of Groningen, Groningen (van der Does, et al., 1998) |
| His secY<sub>M15</sub> EG | P0AGA2<br>P0AG96<br>P0AG99 | pIMBB602 | pET610 | (Karamanou et al., 2008) |
| His secAcys-(V280C/L464C/L187A)secA(L187A) | P10408P10408 | pLMB1910pLMB1860 | pET16bpET5a | The mutation L187A was introduced in pLMB1819 using primer pairs X403-X901secA N31-898 (L187A) fragment from pIMBB680 (His SecA-L187A) was inserted in pLMB0081 after NcoI digestion |
| His secA N6-901 (L187A) His secAcys-(V280C/L464C/L187A) | P10408P10408 | pIMBB680pLMB1910 | pET5apET16b | The mutation L187A was introduced in pIMBB7 using primer pairs X403-X901The mutation L187A was introduced in pLMB1819 using primer pairs X403-X901 |

#### List of Primers

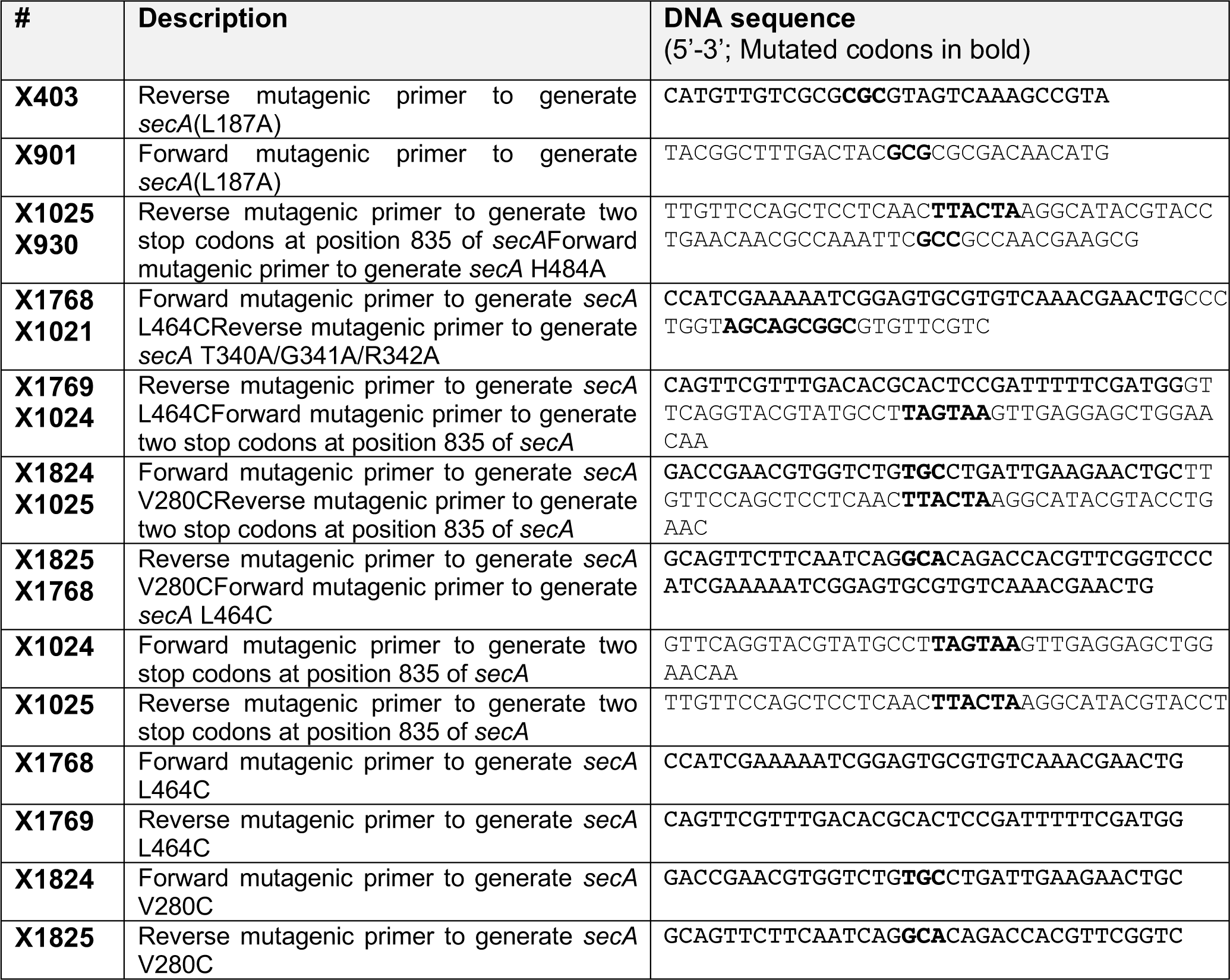
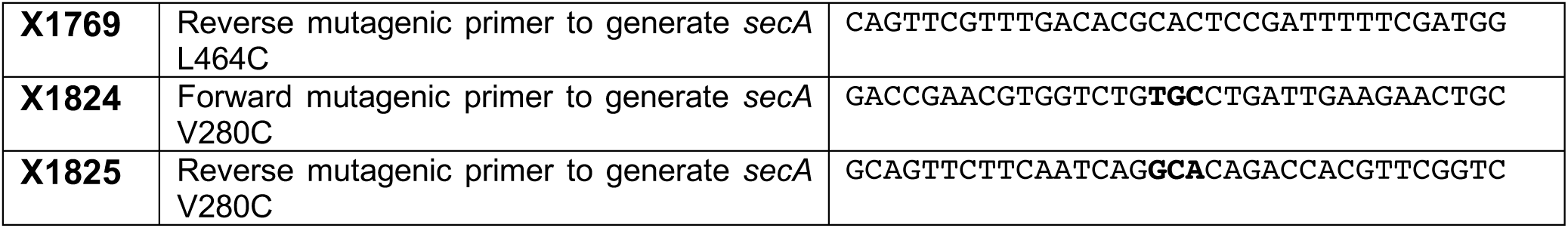

